# PheGWAS: A new dimension to visualize GWAS across multiple phenotypes

**DOI:** 10.1101/694794

**Authors:** Gittu George, Sushrima Gan, Yu Huang, Philip Appleby, A.S. Nar, Radha Venkatesan, Viswanathan Mohan, Colin N.A Palmer, Alex S.F Doney

**Affiliations:** NIHR Global Heath Research Unit on Global Diabetes Outcomes Research, Division of Population Health and Genomics, University of Dundee, Ninewells Hospital and Medical School, Dundee, UK

**Author notes:** These authors contributed equally to the work (Sushrima Gan, Yu Huang).

## Abstract

**Motivation:** PheGWAS was developed to enhance exploration of phenome-wide pleiotropy at the genome-wide level through the efficient generation of a dynamic visualization combining Manhattan plots from GWAS with PheWAS to create a three-dimensional “landscape”. Pleiotropy in sub-surface GWAS significance strata can be explored in a sectional view plotted within user defined levels. Further complexity reduction is achieved by confining to a single chromosomal section. Comprehensive genomic and phenomic coordinates can be displayed.

**Results:** PheGWAS is demonstrated using summary data from Global Lipids Genetics Consortium (GLGC) GWAS across multiple lipid traits. For single and multiple traits PheGWAS highlighted all eight-eight and sixty-nine loci respectively. Further, the genes and SNPs reported in GLGC were identified using additional functions implemented within PheGWAS. Not only is PheGWAS capable of identifying independent signals but also provide insights to local genetic correlation (verified using HESS) and in identifying the potential regions that share causal variants across phenotypes (verified using colocalization tests).

**Availability and Implementation:** The PheGWAS software and code are freely available at (https://github.com/georgeg0/PheGWAS).

## 1 INTRODUCTION

The potential of personalized medicine has evolved extensively in the last decade with the development of genome-wide association studies (GWAS), which is a powerful method for exploring the genetic architecture underlying diseases and traits affecting humans. Large Bioresources like UK Biobank (https://www.ukbiobank.ac.uk) and eMERGE (https://emerge.mc.vanderbilt.edu) that combine genomic data with electronic medical records of study participants provide opportunities to perform GWAS across many different diseases and traits. Systemically analyzing the enormous volume of data produced in these studies is one of the most significant issues at present. The ability to visualize complex data can significantly enhance its exploration and understanding (Li *et al.*, 2012). Applying this to the exploration of many genetic variants over many diseases demands data visualization tools which present the data in an intuitive way that is also capable of efficiently handling very large volumes of data.

The Manhattan plot is the most readily available and established way to visualize GWAS and provides instant appreciation of the underlying genetic structure of the disease or trait being studied. It comprises a scatter plot of the positions of the SNPs along each autosomal chromosome on the x-axis and the y-axis corresponding to the significance of the association (expressed as −log_10_(*p*)) with the particular phenotype in question. In spite of its ubiquitous use in GWAS only the very significant loci can be visualized by this static representation although in this context this is not a limitation as the aim is to identify only the top-most significant SNPs. In order to appreciate underlying deeper structure with weaker associations tools like qqman (D. Turner, 2018) are required. Regional GWAS are also offered by LocusTrack (Cuellar-Partida *et al.*, 2015) which is another tool which combines the features of LocusZoom (Pruim *et al.*, 2010) and SNAP plot (Johnson *et al.*, 2008) and allows to choose between plotting the *p* values or linkage disequilibrium (LD) on the y-axis.

All such GWAS data visualizations are for “many variant - one phenotype” studies. However, when a researcher is interested in pleiotropy (Gratten and Visscher, 2016) and therefore requires to assess if particular variants are associated across a group of phenotypes PheWAS is undertaken which is the reverse of GWAS and considers the “one variant-many phenotypes”, providing a mechanism to detect pleiotropy (Roden *et al.*, 2010). Software like PheWAS-view (Pendergrass *et al.*, 2012) and R-PheWAS (Carroll *et al.*, 2014) have been developed to visualize and summarize PheWAS results at both individual and larger population level, enabling the exploration of pleiotropy for single SNPs (Pendergrass *et al.*, 2012).

We describe a combination of these approaches in a “many variants-many phenotypes” scenario (Fig 1) to enable visualization of genome-wide data plots across a number of phenotypes as a three-dimensional landscape (Fig 2). This approach which we refer to as PheGWAS might assist in understanding or exploring pleiotropy at a genome wide scale (Heinrich *et al.*, 2012).

**Fig 1:**
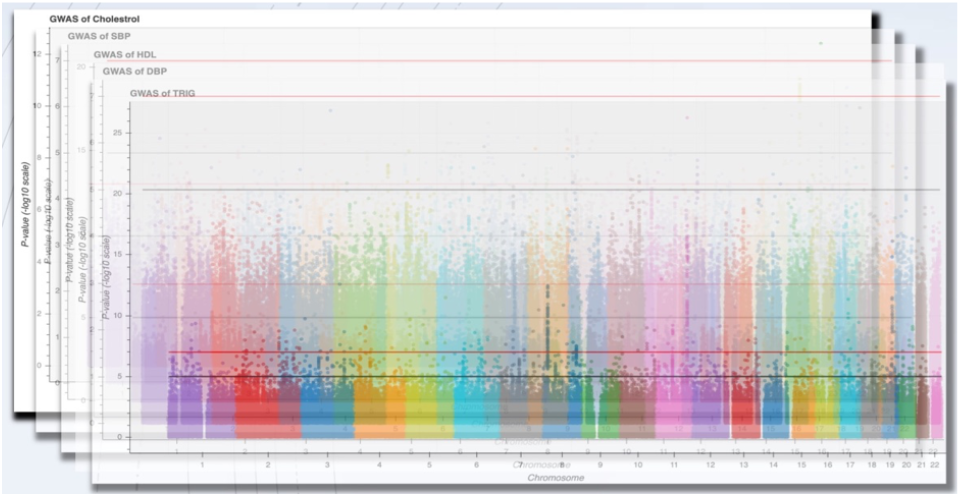
Visualization of the foundational backbone of the PheGWAS concept

**Fig 2:**
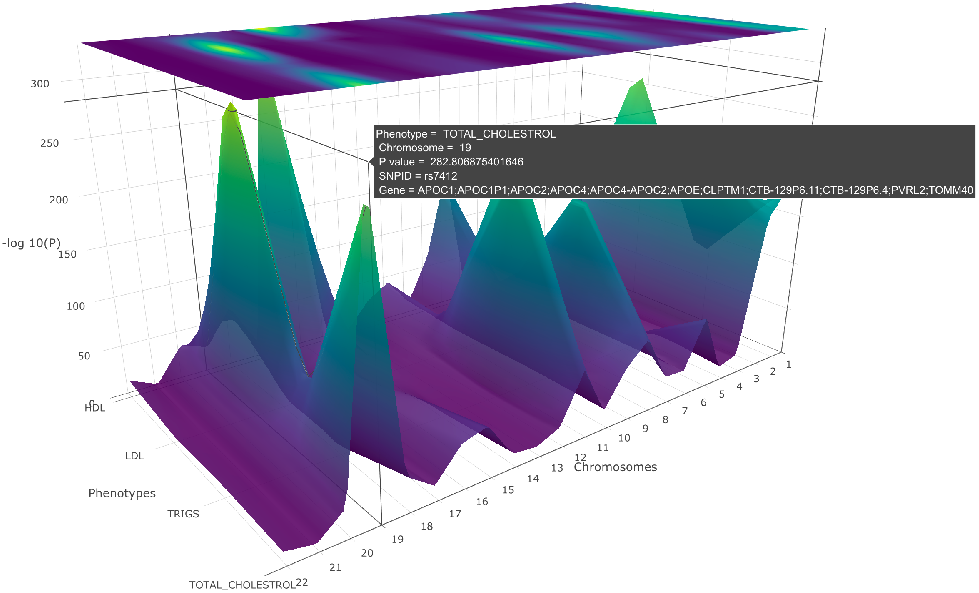
A PheGWAS graph for the phenotypes, SBP, DBP, HDL, Triglycerides and Cholesterol illustrating the interactive landscape. Rotating the x-axis of the graph will allow a clear picture of the

## 2 METHODS

### 2.1 PheGWAS analysis

PheGWAS allows dynamic interactive three-dimensional visualization and exploration of a genome-wide by phenome-wide landscape broadly at two levels – the entire genome level and by single chromosome level. Both of these levels can be explored at any user configurable significance level in a “sectional view”. This is achieved by selecting a significance interval on the y-axis and displaying the peaks in a substratum of the landscape within the selected threshold (Fig 4).

PheGWAS implements orbital rotation and turntable-rotation. Furthermore, it also provides a pan feature to enable an aligned display. Turntable rotation of the x-axis brings us to the heatmap (Fig 3) which is the projection of the surface into the coordinate planes perpendicular to the z-axis in the three-dimensional space. The heatmap is highlighted according to the corresponding y-axis value (i.e., −log_10_(*p*)) and provides a flyover view of the landscape across all chromosomal strips.

**Fig 3:**
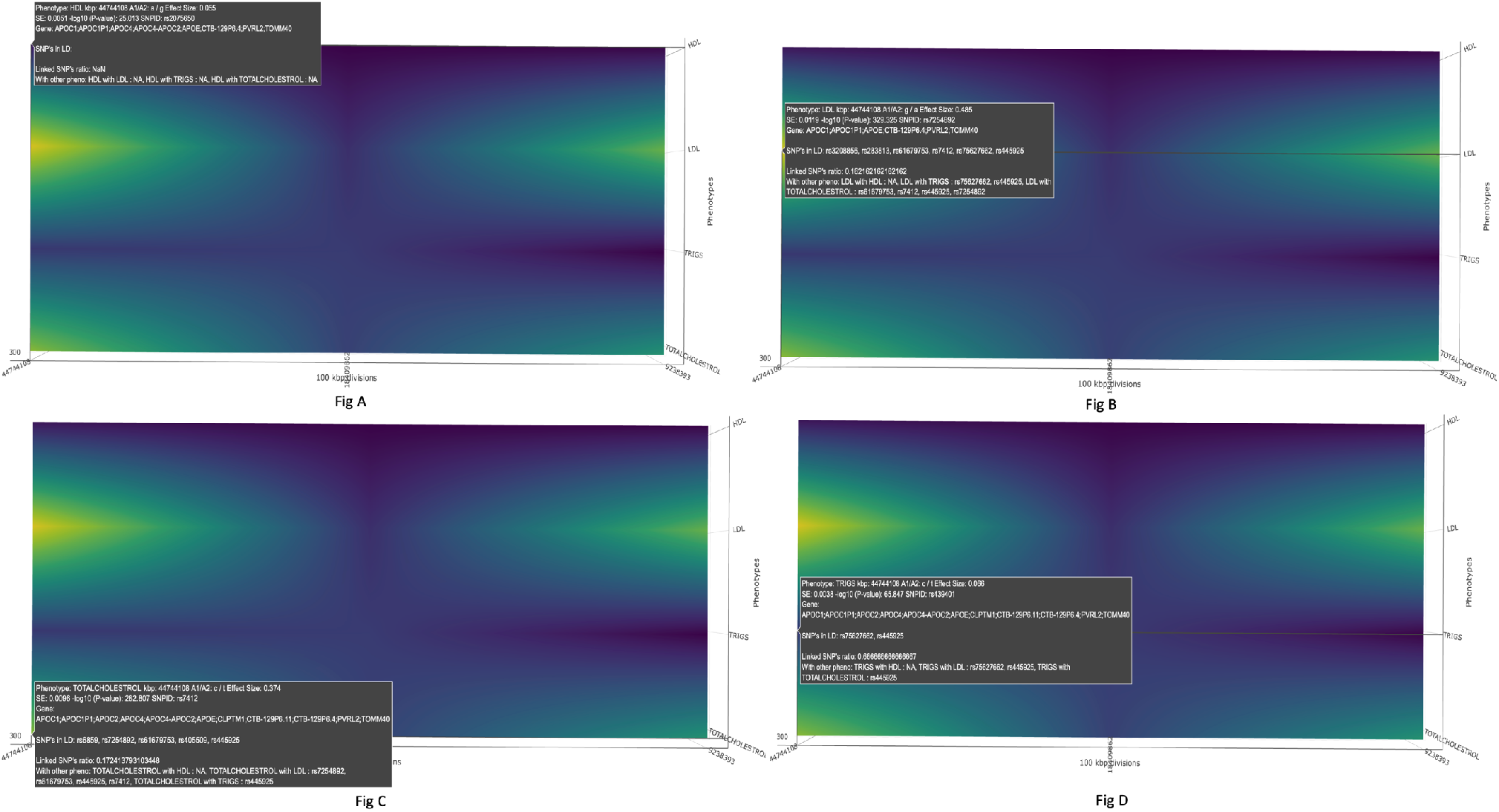
An illustration of the heatmap produced by PheGWAS (single chromosomal view for 19th chromosome). The highlighted regions represent the SNPs with significant −log_10_*(p).* This helps the user to decide which all chromosomes will be selected for the individual level chromosomal view. a) rs2075650 found to be significantly associated (p value: 9.7e-26) with HDL (SNPS in LD: NA) b) rs7254892 found to be significantly associated (p value: 1.6e-320) with LDL (SNPS in LD: rs3208856, rs283813, rs61679753, rs7412, rs75627662, rs445925) c) rs7412 found to be significantly associated (p value: 1.6e-283) with TOTAL CHOLESTROL (SNPS in LD: rs6859, rs7254892, rs61679753, rs405509, rs445925) d) rs439401 found to be significantly associated (p value: 1.4e-66) with TRIGS (SNPS in LD: rs75627662, rs445925)

**Fig 4:**
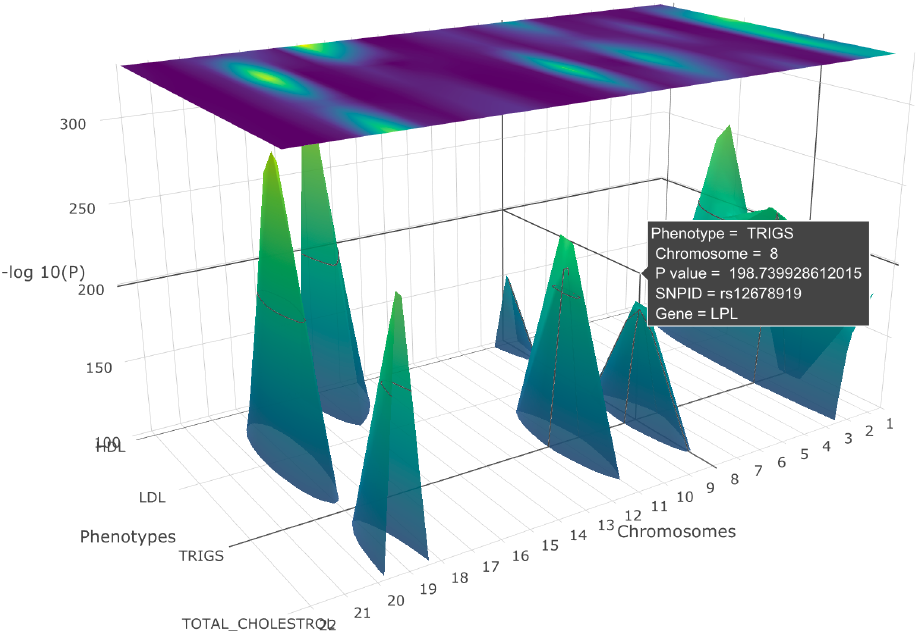
A PheGWAS plot for the phenotypes, SBP, DBP, HDL, Triglycerides and Cholesterol with a sectional view of −log_10_*(p)* greater than 100 (entire genome level)

#### 2.1.1 Entire genome level

At the entire genome level, PheGWAS provides an overview of the entire landscape (Fig 2). Here, as in a conventional Manhattan plot the x-axis represents the autosomal chromosomes (i.e., chromosomes 1-22) and the y-axis represents the −log_10_(*p*) of the GWAS summary statistics the additional z-axis represents the range of phenotypes. To de-clutter the view, considering that there may be multiple peaks in each chromosome, only the maximum −log_10_(*p*) is selected as is the case for each user selectable substrata (see below).

Each of the phenotypes are overlaid simultaneously on the same graph giving rise to the “see-through” landscape topology. Axis grid lines appear spontaneously as a cursor is moved over the landscape to show a precise position of the SNPs. At exact SNP positions, a dialog box appears showing the phenotype, chromosome, SNP ID, −log_10_*(p)*, effect size, standard error, allele 1, allele 2, locus and the gene symbol. This provides a quick orientation point on the PheGWAS landscape.

#### 2.1.2 Single chromosomal plot

From this entire genome overview the user can select a single chromosome for more granular exploration (Fig 5). Here the x-axis corresponds to base pair interval segments (see below) along the selected chromosome. As for the entire genome level, the y and z axis correspond to the −log_10_(*p*) of the GWAS summary statistics and the phenotypes respectively. Several background processes are required to produce the single chromosome plot.

**Fig 5:**
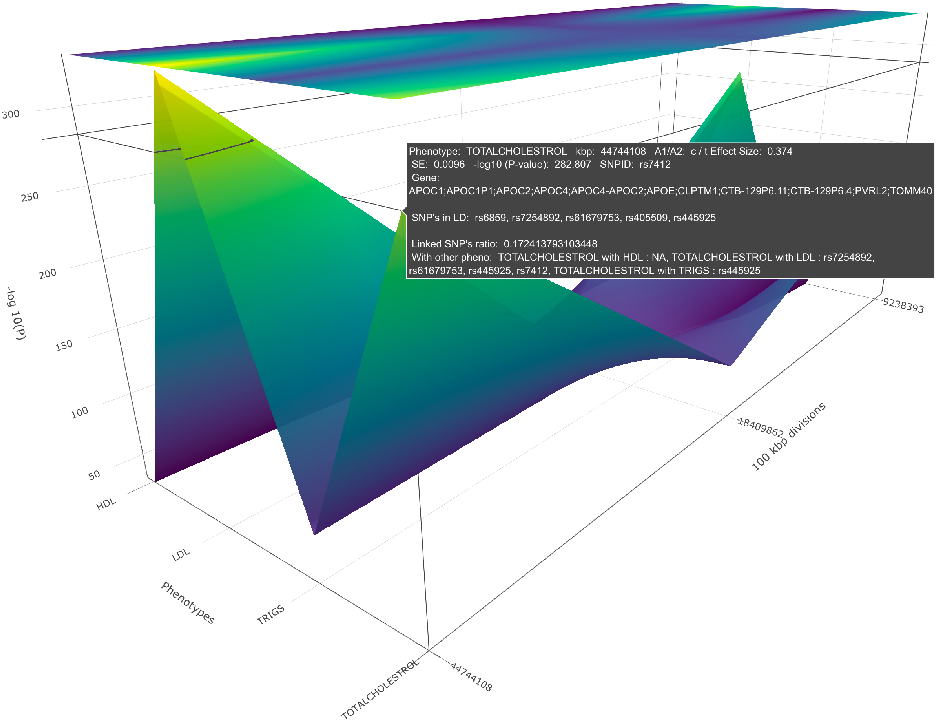
A PheGWAS plot for the sectional view of a single chromosome (19th chromosome), produced by plotting the SNPs above a certain threshold of significant values of phenotypes,

The length of the chromosome can be divided into equal base pair interval segments predefined by the user (default 1M base pairs), giving rise to a systematic order of groups of SNPs. Alternatively it can be divided based on LD blocks (Berisa and Pickrell, 2016) defined by the population (e.g. Asian, European, or African), where the input is the conventional ‘*bed*’ file that has chromosome name and the start and end position of the block. This provides the opportunity of discriminating peaks that represent pleiotropy across phenotypes or simply constitute discrete separate signals.

A base pair interval segment in a single chromosome is selected for display only if there are one or more peaks in that segment along the z-axis that have a −log_10_(*p*) greater than 6 (minimum significant threshold). Similar to the sectional view for the whole genome, a sectional view with a minimum significant threshold can be chosen for the single chromosomal plot. Within a selected base pair interval segment, only the highest peak for each phenotype (z-axis) is plotted, any base pair interval segment which has no peak greater than the selected minimum significant threshold is omitted from the plot making. Here again, the axis grid lines help to spot a more precise location of the SNP.

##### 2.1.2.1 Modules implemented in single chromosomal view

To facilitate in depth exploration of the PheGWAS landscape several example modules have been implemented within the single chromosome view. It is anticipated that these will be further developed and added to by users.

###### Gene View Heat Map

Researchers might find the visualization of SNPs challenging if there is a large variation in the −log_10_(*p*). This is because a particular SNP with the highest −log_10_(*p*) and its corresponding phenotype will be highlighted in the heatmap at the top of the heatmap scale. As a consequence of the available scale range an adjacent −log_10_*(p)* may still be significant but may not be highlighted in the heatmap potentially concealing the association from the researcher. To counter this, PheGWAS provides a different heatmap view by identifying identical genes across phenotypes within a user customized base pair group. This heatmap highlights phenotypes that share variants within the same gene locus and highlights all the SNPs above the pre-selected threshold −log_10_(*p*) with uniform brightness.

###### Effect Size Plot

PheGWAS allows the user to view the relative size of the biological effect by providing effect size (Beta or OR) on the y-axis in this plot rather than −log_10_*(p).* The heatmap continues to be highlighted according to the −log_10_*(p)* allowing the user to view both statistics simultaneously (Fig 6).

**Fig 6:**
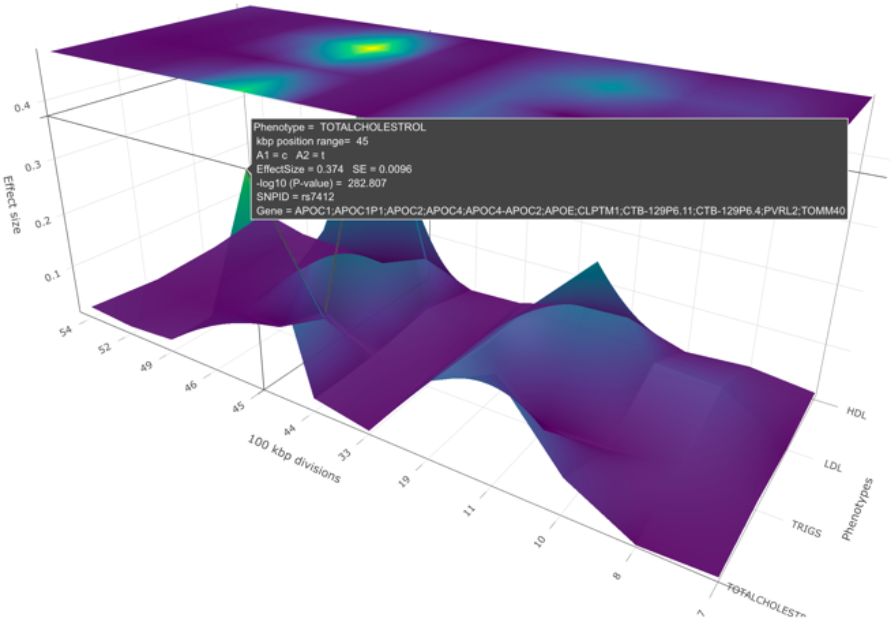
A PheGWAS effect size plot for the sectional view of a single chromosome (19th chromosome), produced by plotting the SNPs above a certain threshold of significant values of phenotypes, SBP, DBP, HDL, Triglycerides and Cholesterol

###### Linkage disequilibrium (LD) block-based chromosome segmentation

In a base pair interval segment PheGWAS can identify SNPs in LD with the most significant SNP being plotted. Here the LD threshold and population used to derive the LD are user customizable. An option to find the mutual LD SNPs between the phenotypes has been featured, to assist identifying SNPs that are in LD shared between phenotypes. This is displayed in the pop-up window.

In addition, PheGWAS identifies the number of SNPs which are in LD with the SNP with highest −log_10_*(p)* and the total number of SNPs above the set threshold. Finally, the ratio of these two gives the proportion of linked SNPs with genome-wide significance which is displayed in the pop-up along with the other information.

While selecting the highest peak declutters the image for viewing, the LD module is helpful to further verify information such as whether a signal is independent or in identifying false signals.

###### SNP thinning based on LD functionality

This function allows the user to capture the independent signals or the signals which are buried inside a base pair interval segment. By default, PheGWAS displays the SNP with the highest significance. Following this, all the SNPs which are in LD with this SNP are removed. This is similar to clumping (http://zzz.bwh.harvard.edu/plink/clump.shtml) but without the requirement of having individual level data. The user can perform this process recursively and specify the depth of moving downwards for independent signals within a base pair interval segment.

###### Export regional data for further analysis

If a region is found to be of interest, then the summary statistics of this region can be exclusively exported to undertake further analysis such as identifying colocalized signals with external tools (see section 3.3).

## 3 IMPLEMENTATION

The PheGWAS code has been scripted in R. The code is wrapped as a package executed in two phases. The first phase (executed once) accepts the GWAS summary files as R data-frames to format it into a single data-frame that can be accepted in the second phase. By default, in the second phase, PheGWAS generates an interactive 3D plot for all chromosomes. To plot an individual chromosome, the chromosome number must first be provided. Similarly, for the sectional view, a −log_10_(*p*) interval is required. PheGWAS plots are rendered using a package called ‘plotly’ (https://plotly-r.com). There is an option to save the plots as an interactive HTML webpage or a static diagram. It is recommended to save this as interactive HTML as this gives the full power to the user to explore and demonstrate the data within the visualization.

SNP ID and genome location data within the GWAS file is mapped to the respective GENE region that it falls into. This mapping is implemented using the *BioMart* R package, which gives the associated gene for each SNP ID in the GWAS summary file.

PheGWAS connects to ensemble (https://www.ensembl.org) to get the LD values between the top significant SNP and all other variants within the same base pair interval segment. SNPs reaching a certain threshold (default r^2^ = 0.75 and d’ = 0.75) are shown in the pop-up.

For the demonstration provided in this paper (each summary statistics files having ~ 2.5 million SNPs), the first phase of PheGWAS takes approximately 284,000 milliseconds and second phase takes approximately 400 milliseconds to complete (tested against running function in a server with 8 GB RAM).

### 3.1 PheGWAS applied example: Using Global Lipids Genetics Consortium (GLGC) summary statistics file

To demonstrate and validate PheGWAS, we used summary data from GLGC (Grallert *et al.*, 2013). GLGC examined 188,577 individuals to identify new loci and refine estimates for previously known loci influencing plasma levels of low-density lipoprotein (LDL), high-density lipoprotein (HDL) cholesterol, triglycerides and total cholesterol. This revealed eighty-eight SNPs associated with single traits and the sixty-nine SNPs associated with multiple traits. For the current PheGWAS analysis, a cut-off −log_10_(*p*) of 6.5 and base pair division value of 1M base pair was used.

The 19^th^ chromosome was selected to showcase a detailed PheGWAS analysis and the possible inferences from it. By viewing the heatmap of the z/x plane, the pattern of genomic significant region across the whole chromosome for multiple traits is demonstrated.

#### 3.1.1 Preparing the summary statistics file

In order to visualize the GLGC data on a PheGWAS plot it was necessary to first map SNPs to respective genes using the BioMart package (http://bioconductor.org/packages/release/bioc/html/biomaRt.html). Because this process does not allow for customized mapping windows of SNPs to genes, to keep it in line with GLGC, a 100K base pair interval window was used to manually map genes. Files from two sources were used for mapping. One from UCSC genome-mysql.soe.ucsc.edu ftp server for SNP rsid’s with respective chromosomes start and end position and the other from the Genome Reference Consortium Human Build 37 (GRCh37), for the genes to map to the chromosome position. These were used to map the gene names to SNPs, using bedmap (https://bedops.readthedocs.io/) to look for SNPs whose positions overlap with the genes in the 100K base pair window.

### 3.2 PheGWAS validation

Validation was carried out at the chromosomal level for each of the twenty-two chromosomes. For single and multiple traits, there were three stages: -

i. Overall significant SNP Display – Whether the same significant SNPs region as reported by GLGC were displayed in the PheGWAS heatmap.
ii. Corresponding Gene Symbol display – This determined whether the gene mapping process for the GLGC summary data in PheGWAS had been successful so that the gene symbol associated with a particular SNP in GLGC data is the same as gene indicated by PheGWAS
iii. Identifying SNPs by base pair interval – In this process it was determined whether PheGWAS successfully indicated the same SNP within the same 100kb base pair interval segment as GLGC (chromosomal locus). SNPs highlighted by PheGWAS in a particular base pair interval segment with the lowest *p* value (the default) that were identical to those reported by GLGC were categorized as A SNPs reported by GLGC are in LD (using LD functionality; if r^2^>0.75 and D’>0.75) with the SNPs highlighted by PheGWAS in a particular base pair interval segment were categorized as B. If the SNP reported by GLGC are neither in category A nor in category B, then it is categorized as C and we apply the SNP thinning functionality to see if we can find the buried SNP in the base pair interval segment. The level mentioned in bracket will indicate the depth at which the SNP is found.

### 3.3 Further analyses on data exported from PheGWAS

For the purpose of verifying the local genetic correlation we used a quantitative method -ρ-HESS (Shi *et al.*, 2017) across phenotypes. ρ-HESS analysis was carried out only on TC and LDL as ρ-HESS is only capable considering 2 traits at a time.

If a genomic region is identified as having a shared genetic component across two or more phenotypes, it’s important to know if a potentially causal variant is involved. Usually a colocalization test is done on this region to see if there are colocalized traits and identify any causal variant involved. Available statistical software’s like MOLOC (Giambartolomei *et al.*, 2018) and HyPrColoc (Foley *et al.*, 2019)were used for performing colocalization tests for multiple phenotypes. The base pair region 45 in the 19^th^ chromosome was selected as an example.

## 4 RESULTS

### 4.1 Demonstration of PheGWAS – A Walkthrough

#### 4.1.1 Entire genome level

From the initial entire genome view, for chromosome 19, four genomic peak regions in the interactive visualization were demonstrated corresponding to HDL, LDL and Triglycerides and Total Cholesterol (Fig 2. NB only the largest two peaks are visible in this static rendition).

#### 4.1.2 Single 19^th^ chromosomal level

To locate the exact position of the SNPs in these four peaks the 19^th^ chromosome is selected for the single chromosomal view. Here a view of the significant peaks for each of the base pair regions is available (Fig 3). The cursor is hovered over the heatmap to carry out an inspection of base pair positions.

Further, by viewing the heatmap (Fig 3) of the PheGWAS plot for GLGC traits, several GWAS significant genomic regions were observed. This suggests further scrutiny of the shared genetic architecture in this region.

Using the data export function local genetic correlation across phenotypes LDL and TC was verified using ρ-HESS plot. The peaks that were found in PheGWAS plot (Supplementary Fig 1) were also found in the ρ-HESS plot (Supplementary Fig 2).

The PheGWAS plot indicates that base pair region 45 in the 19^th^ chromosome contains a genetic architecture shared between the phenotypes. The popup window shows the number of SNPs that reached significant threshold in this region and are shared between different traits. (Fig3). This suggests performing a multiple trait colocalization analysis. The export function was used to get the summary statistics of this base pair region which was used to perform multiple traits colocalization analysis. The results are shown (Supplementary Section 1).

### 4.2 Summarizing PheGWAS results using GLGC summary statistics file

For displaying overall significant SNPs from GLGC, the heatmap was able to provide a visual display of all the significant regions identified within GLGC data (Supplementary tables 1a and 2a – column 6). However, this was verified with tables provided by PheGWAS (Supplementary tables 3 - 23).

The corresponding gene symbols identified by PheGWAS were also compared to the ones identified by GLGC. In single traits, seventy-seven genes reported by GLGC within that base pair interval segment were also identified by PheGWAS (Supplementary tables 1a - column 7). In multiple traits, fifty-nine genes reported by GLGC within that base pair interval segment were also identified by PheGWAS (Supplementary tables 2a - column 7). Any minor discrepancies were attributed to different resources used for gene annotation.

For SNP identification by base pair interval, in single traits, sixty-three SNPs reported by GLGC with the lowest (*p* value) within that base pair interval segment were also identified by PheGWAS (Category A, Supplementary tables 1a – column 8). Twenty-two SNPs detected by PheGWAS were in LD (Supplementary tables 1c) with the SNP reported by GLGC (Category B, Supplementary tables 1a– column 8). Three SNPs were found by SNP thinning functionality (Category C, Supplementary tables 1a-column 8). For multiple traits, twenty one SNPs reported by GLGC with the lowest (*p* value) within that base pair interval segment were identified by PheGWAS and were marked as A (Supplementary table 2a – column 8), forty four SNPs detected by PheGWAS were in LD (Supplementary tables 2c) with the SNP reported by GLGC and marked as B (Supplementary table 2a – column 8) and four SNPs were found by SNP thinning functionality and marked as C (Supplementary tables 2a– column 8).

The summary table for single and multiple traits have been provided separately. (Supplementary tables 1b and 2b)

A complete list of the PheGWAS findings of genes and SNPs of all the twenty-two chromosomes is provided (Supplementary Tables 3-23).

## 5 DISCUSSION AND CONCLUSION

PheGWAS creates a new three-dimensional visualization approach for many SNPs against many phenotypes to aid scientific research. Endeavors to visualize results of multiple GWASs on a single plot have been made in the past. Researchers like Wang et al.(Wang *et al.*, 2016) and Hoffman et al.(Hoffmann *et al.*, 2018) have demonstrated visualizing results of more than one GWAS simultaneously to view multiple regions associated with a particular phenotype, by overlapping two Manhattan plots. There are shortcomings of static, non-interactive data visualization. Interactive data visualization permits more comprehensible representation supporting the user to find solutions to distinct scientific problems(Khramtsova and Stranger, 2017). Further developments have been made to make these static plots more interactive for the user by R/qtlcharts(Broman, 2015) and Zbrowse(Ziegler *et al.*, 2015). Visualization of the Manhattan plot becomes an insurmountable challenge when hundreds of thousands of SNPs are plotted. To deal with this existing interactive browsers for visualization of multiple GWAS experiments such as Zbrowse (Ziegler *et al.*, 2015) and Assocplots (Khramtsova and Stranger, 2017) have put a restriction on the sample size by selecting top 5000 and 1000 SNPs respectively.

Unlike the static and non-interactive Manhattan plot, PheGWAS does not plot all existing points but only focuses on SNPs over a certain pre-decided level of significance. The advantages of the PheGWAS plot are many; genetic variants can be browsed over multiple phenotypes in a single user definable plot which can be dynamically manipulated. Also, compared to a PheWAS plot, PheGWAS plots provide the chance to view different regions and traits at the same time allowing immediate identification of pleiotropic effects for multiple loci and multiple phenotypes. This enhances the researcher’s opportunity for subsequent data analysis such as appropriate covariate selection in identifying genetic modifiers or for the construction of various predictive models that estimates the effect of a predictor on the response, in the presence of pleiotropy, like Genetic Instrumental Variable regression.

Using the example of the GLGC summary data it was seen that PheGWAS is highly efficient in identifying SNPs and associated genes with an additional visual landscape representation by providing the researcher a walk-through of the SNP hit exploration. Although the current state of PheGWAS cannot perform any statistical analysis for colocalization it can provide export data of regions of interest within the chromosome view that can be taken further for a colocalization study as demonstrated. This identification process of the region from the PheGWAS plot makes comparison of the association signals visually seamless rather than the traditional method of manually overlaying plots on top of each other (Kanduri *et al.*, 2019).

While our demonstration considered a number of phenotypes PheGWAS can also be used to identify the different genetic architecture for the same trait between cohorts, such as population stratifications based on ethnicities and gender, provided the GWAS summary statistics data is available for each sub-group (in the package the gender data of BMI from giant consortium is provided). It does not, however, put any restriction on the number of sub-groups to be plotted.

Some of the limitations of PheGWAS were identified – it does not consider sample size variation, does not integrate by weighing the p-values, and insights towards the local genetic correlations can’t explain the negative correlation between the traits and the region. PheGWAS is an ongoing project in which efforts are being made to implement these techniques for making it more appropriate for comprehensive gene-disease association analysis.

## 6 FUTURE IMPLEMENTATIONS

In PheGWAS, at each level, the plots allow investigation of each chromosome in greater detail than the previous level. In this report we have demonstrated the highest level of visualization of all chromosomes, sectional visualization of significance strata and the single chromosomal view. We envisage a progression to furthermore detailed views being developed by the user community on the basis of the shared open source software in GitHub. Such as creating a three-dimensional scatter plot for a specific base pair group and significance interval that expand the two-dimensional locus-specific plots for single phenotype SNP stack provided by LocusZoom (Pruim *et al.*, 2010) into multi-phenotype three dimensions. We also envisage future incorporation of additional information in SNP annotations by connecting to external databases such as DisGeNET (http://www.disgenet.org) for gathering published research about associations of gene with any other diseases. Functionality for adding expression quantitative trait loci (eQTL) and methylation quantitative trait loci (mQTL) to the plot will help to understand the molecular alterations within a selected region in PheGWAS. The provision of information about the SNPs in LD as well as a measure of pleiotropic effects, if present, similar to the one provided by ShinyGPA (Kortemeier *et al.*, 2018) we believe would also be useful.

We are considering a future implementation of RShiny (an R package to build interactive web applications) to provide greater flexibility for interacting when passing parameters to PheGWAS. The Shiny program could be hosted in any local server and be used as an interactive way to pass the parameters for a PheGWAS. Interactive nature gives the user the ability to upload the GWAS summary files to the web interface and then perform the PheGWAS on the entire chromosome or any chromosomes from the dropdown menu. The threshold to use for the sectional view can also be set according to the user preference.

## Acknowledgements

We would like to thank the Health Informatics Centre (HIC), the principal investigators and our colleagues in the INSPIRED project.

