## Supplementary File for "PheGWAS: A new dimension to visualize GWAS across multiple phenotypes"

### TABLES:

**Supplementary Table 1a:** Comparative analysis loci identified by GLGC and PheGWAS results for association of loci with single traits

| Nearest Gene | SNP ID | Chromosome | Position MB | Trait | In PheGWAS | Gene Match | Identifying GLGC SNP's |
|---|---|---|---|---|---|---|---|
| *EVI5* | rs7515577 | 1 | 93.01 | TC | DETECTED | M | B |
| *ZNF648* | rs1689800 | 1 | 182.17 | HDL | DETECTED | NM | B |
| *PABPC4* | rs4660293 | 1 | 40.03 | HDL | DETECTED | M | A |
| *ASAP3* | rs1077514 | 1 | 23.77 | TC | DETECTED | M | A |
| *ANXA9* | rs267733 | 1 | 150.96 | LDL | DETECTED | M | A |
| *RRNAD1* | rs12145743 | 1 | 156.70 | HDL | DETECTED | M | A |
| *C1orf220* | rs4650994 | 1 | 178.52 | HDL | DETECTED | M | A |
| *RAB3GAP1* | rs7570971 | 2 | 135.84 | TC | DETECTED | M | B |
| *COBLL1* | rs12328675 | 2 | 165.54 | HDL | DETECTED | M | B |
| *EHBP1* | rs2710642 | 2 | 63.15 | LDL | DETECTED | M | A |
| *ABCB11* | rs2287623 | 2 | 169.83 | TC | DETECTED | M | A |
| *FAM117B* | rs11694172 | 2 | 203.53 | TC | DETECTED | M | A |
| *CPS1* | rs1047891 | 2 | 211.54 | HDL | DETECTED | M | A |
| *FN1* | rs1250229 | 2 | 216.30 | LDL | DETECTED | M | A |
| *RAF1* | rs2290159 | 3 | 12.63 | TC | DETECTED | NM | C (level 3) |
| *MSL2L1* | rs645040 | 3 | 135.93 | TG | DETECTED | M | A |
| *ATG7* | rs2606736 | 3 | 11.40 | HDL | DETECTED | M | A |
| *SETD2* | rs2290547 | 3 | 47.06 | HDL | DETECTED | M | A |
| *RBM5* | rs2013208 | 3 | 50.13 | HDL | DETECTED | M | A |
| *STAB1* | rs13326165 | 3 | 52.53 | HDL | DETECTED | M | A |
| *PXK* | rs13315871 | 3 | 58.38 | TC | DETECTED | M | A |
| *GSK3B* | rs6805251 | 3 | 119.56 | HDL | DETECTED | M | A |
| *KLHL8* | rs442177 | 4 | 88.03 | TG | DETECTED | NM | A |
| *SLC39A8* | rs13107325 | 4 | 103.19 | HDL | DETECTED | M | A |
| *C4orf52* | rs10019888 | 4 | 26.06 | HDL | DETECTED | NA | A |
| *FAM13A* | rs3822072 | 4 | 89.74 | HDL | DETECTED | M | A |
| *ADH5* | rs2602836 | 4 | 100.01 | HDL | DETECTED | M | A |
| *ARL15* | rs6450176 | 5 | 53.30 | HDL | DETECTED | M | A |
| *MAP3K1* | rs9686661 | 5 | 55.86 | TG | DETECTED | NM | A |
| *CITED2* | rs605066 | 6 | 139.83 | HDL | DETECTED | NM | B |
| *C6orf106* | rs2814982 | 6 | 34.55 | TC | DETECTED | M | A |
| *KCNK17* | rs2758886 | 6 | 39.25 | TC | DETECTED | M | A |
| *HBS1L* | rs9376090 | 6 | 135.41 | TC | DETECTED | M | A |
| *TYW1B* | rs13238203 | 7 | 72.13 | TG | DETECTED | NM | B |
| *KLF14* | rs4731702 | 7 | 130.43 | HDL | DETECTED | M | B |
| *GPR146* | rs1997243 | 7 | 1.08 | TC | DETECTED | M | A |
| *DAGLB* | rs702485 | 7 | 6.45 | HDL | DETECTED | M | A |
| *SNX13* | rs4142995 | 7 | 17.92 | HDL | DETECTED | M | A |
| *IKZF1* | rs4917014 | 7 | 50.31 | HDL | DETECTED | M | A |
| *MET* | rs38855 | 7 | 116.36 | TG | DETECTED | M | A |
| *TMEM176A* | rs17173637 | 7 | 150.53 | HDL | DETECTED | M | A |
| *PINX1* | rs11776767 | 8 | 10.68 | TG | DETECTED | M | B |
| *TRPS1* | rs2293889 | 8 | 116.60 | HDL | DETECTED | M | A |
| *JMJD1C* | rs10761731 | 10 | 65.03 | TG | DETECTED | M | B |
| *CYP26A1* | rs2068888 | 10 | 94.84 | TG | DETECTED | M | A |
| *AKR1C4* | rs1832007 | 10 | 5.25 | TG | DETECTED | M | A |
| *VIM* | rs10904908 | 10 | 17.26 | TC | DETECTED | M | A |
| *SPTY2D1* | rs10128711 | 11 | 18.63 | TC | DETECTED | M | B |
| *AMPD3* | rs2923084 | 11 | 10.39 | HDL | DETECTED | M | A |
| *LRP4* | rs3136441 | 11 | 46.74 | HDL | DETECTED | NM | A |
| *OR4C46* | rs11246602 | 11 | 51.51 | HDL | DETECTED | M | A |
| *PCNXL3* | rs12801636 | 11 | 65.39 | HDL | DETECTED | M | A |
| *MOGAT2* | rs499974 | 11 | 75.46 | HDL | DETECTED | M | A |
| *PHLDB1* | rs11603023 | 11 | 118.49 | TC | DETECTED | M | A |
| *PDE3A* | rs7134375 | 12 | 20.47 | HDL | DETECTED | NA | B |
| *MVK* | rs7134594 | 12 | 110.00 | HDL | DETECTED | M | B |
| *SBNO1* | rs4759375 | 12 | 123.80 | HDL | DETECTED | M | C (level 1) |
| *SCARB1* | rs838880 | 12 | 125.26 | HDL | DETECTED | M | B |
| *PHC1* | rs4883201 | 12 | 9.08 | TC | DETECTED | M | A |
| *BRCA2* | rs4942486 | 13 | 32.95 | LDL | DETECTED | M | A |
| *NYNRIN* | rs8017377 | 14 | 24.88 | LDL | DETECTED | M | A |
| *ZBTB42* | rs4983559 | 14 | 105.28 | HDL | DETECTED | M | A |
| *FRMD5* | rs2929282 | 15 | 44.25 | TG | DETECTED | M | B |
| *CAPN3* | rs2412710 | 15 | 42.68 | TG | DETECTED | M | A |
| *LACTB* | rs2652834 | 15 | 63.40 | HDL | DETECTED | M | A |
| *CTF1* | rs11649653 | 16 | 30.92 | TG | DETECTED | M | A |
| *LCAT* | rs16942887 | 16 | 67.93 | HDL | DETECTED | M | A |
| *CMIP* | rs2925979 | 16 | 81.53 | HDL | DETECTED | M | A |
| *PDXDC1* | rs3198697 | 16 | 15.13 | TG | DETECTED | M | A |
| *STARD3* | rs11869286 | 17 | 37.81 | HDL | DETECTED | M | B |
| *PGS1* | rs4129767 | 17 | 76.40 | HDL | DETECTED | M | B |
| *ABCA8* | rs4148008 | 17 | 66.88 | HDL | DETECTED | M | A |
| *MPP3* | rs8077889 | 17 | 41.88 | TG | DETECTED | M | A |
| *APOH* | rs1801689 | 17 | 64.21 | LDL | DETECTED | M | A |
| *MC4R* | rs12967135 | 18 | 57.85 | HDL | DETECTED | NM | B |
| *ANGPTL4* | rs7255436 | 19 | 8.43 | HDL | DETECTED | M | B |
| *FLJ36070* | rs492602 | 19 | 49.21 | TC | DETECTED | NM | B |
| *LILRA3* | rs386000 | 19 | 54.79 | HDL | DETECTED | M | B |
| *ANGPTL8* | rs737337 | 19 | 11.35 | HDL | DETECTED | NM | A |
| *INSR* | rs7248104 | 19 | 7.22 | TG | DETECTED | M | A |
| *FPR3* | rs17695224 | 19 | 52.32 | HDL | DETECTED | M | A |
| *ERGIC3* | rs2277862 | 20 | 34.15 | TC | DETECTED | M | B |
| *SPTLC3* | rs364585 | 20 | 12.96 | LDL | DETECTED | M | A |
| *SNX5* | rs2328223 | 20 | 17.85 | LDL | DETECTED | NM | A |
| *UBE2L3* | rs181362 | 22 | 21.93 | HDL | DETECTED | M | C (level 1) |
| *PLA2G6* | rs5756931 | 22 | 38.55 | TG | DETECTED | M | B |
| *MTMR3* | rs5763662 | 22 | 30.38 | LDL | DETECTED | M | A |
| *TOM1* | rs138777 | 22 | 35.71 | TC | DETECTED | M | A |

In this table, the columns in pheGWAS, gene match and SNP match corresponds to the PheGWAS results we obtained.

Match (M); if the gene symbol in a particular PheGWAS base pair interval segment is the same trait as reported in the GLGC
No Match (NM); if the gene is not identified by PheGWAS.

A- SNP identical with GLGC and PheGWAS
B – GLGC SNP in LD with SNP identified by PheGWAS
C – SNP identified by SNP thinning

**Supplementary Table 1b:** Summary of comparative analysis of loci identified by GLGC and PheGWAS results for association of loci with single traits

| | |
|---|---|
| Loci identified by GLGC | 88 |
| Total Loci Detected by PheGWAS | 88 |
| Gene Match | 77 |
| Gene No Match | 11 |
| Gene not mapped | 2 |
| SNP's in category A | 63 |
| SNP's in category B | 22 |
| SNP's in category C | 3 |

**Supplementary Table 1c:** LD Score between SNPs in GLGC and PheGWAS in single traits

| Chromosome | Position MB | Trait | SNP Match | SNP ID | SNP ID phegwas | d' | r2 |
|---|---|---|---|---|---|---|---|
| 1 | 93.01 | TC | M | rs7515577 | rs1556562 | 1 | 1 |
| 1 | 182.17 | HDL | M | rs1689800 | rs1689797 | 0.99 | 0.89 |
| 2 | 135.84 | TC | M | rs7570971 | rs6730157 | 0.99 | 0.99 |
| 2 | 165.54 | HDL | M | rs12328675 | rs7607980 | 1 | 0.98 |
| 3 | 12.63 | TC | NM | rs2290159 | rs7616006 | 0.33 | 0.04 |
| 6 | 139.83 | HDL | NM | rs605066 | rs3861397 | 0.99 | 0.69 |
| 7 | 72.13 | TG | NM | rs13238203 | rs11974409 | 0.76 | 0.08 |
| 7 | 130.43 | HDL | M | rs4731702 | rs11765979 | 0.99 | 0.99 |
| 8 | 10.68 | TG | NM | rs11776767 | rs6995541 | 0.97 | 0.62 |
| 10 | 65.03 | TG | M | rs10761731 | rs10761762 | 1 | 0.8 |
| 11 | 18.63 | TC | M | rs10128711 | rs10832962 | 1 | 1 |
| 12 | 20.47 | HDL | M | rs7134375 | rs11045163 | 0.95 | 0.87 |
| 12 | 110.00 | HDL | M | rs7134594 | rs10850380 | 1 | 1 |
| 12 | 123.80 | HDL | NM | rs4759375 | rs2454722 | 0.2 | 0.2 |
| 12 | 125.26 | HDL | M | rs838880 | rs838876 | 0.94 | 0.84 |
| 15 | 44.25 | TG | M | rs2929282 | rs16948098 | 0.97 | 0.84 |
| 17 | 37.81 | HDL | M | rs11869286 | rs113612868 | 1 | 1 |
| 17 | 76.40 | HDL | NM | rs4129767 | rs4969178 | 0.94 | 0.57 |
| 18 | 57.85 | HDL | M | rs12967135 | rs6567160 | 1 | 1 |
| 19 | 8.43 | HDL | M | rs7255436 | rs2278236 | 1 | 0.99 |
| 19 | 49.21 | TC | M | rs492602 | rs516246 | 0.99 | 0.99 |
| 19 | 54.79 | HDL | M | rs386000 | rs103294 | 0.99 | 0.9 |
| 20 | 34.15 | TC | NM | rs2277862 | rs72664396 | 0.32 | 0.0002 |
| 22 | 21.93 | HDL | NM | rs181362 | rs113359481 | NA | NA |
| 22 | 38.55 | TG | NM | rs5756931 | rs3761445 | 0.73 | 0.49 |

**Supplementary Table 2a:** Comparative analysis of loci identified by GLGC and PheGWAS results for association of loci with multiple traits

| Nearest Gene | SNP ID | Chromosome | Position MB | Multiple Traits | In PheGWAS | Gene M | Identifying GLGC SNP's |
|---|---|---|---|---|---|---|---|
| *LDLRAP1* | rs12027135 | 1 | 25.78 | TC,LDL | DETECTED | NM | B |
| *PCSK9* | rs2479409 | 1 | 55.50 | LDL,TC | DETECTED | M | B |
| *ANGPTL3* | rs2131925 | 1 | 63.03 | TG,LDL,TC | DETECTED | M | B |
| *SORT1* | rs629301 | 1 | 109.82 | LDL,TC | DETECTED | M | B |
| *MOSC1* | rs2642442 | 1 | 220.97 | TC,LDL | DETECTED | NM | B |
| *GALNT2* | rs4846914 | 1 | 230.30 | HDL,TG | DETECTED | M | B |
| *IRF2BP2* | rs514230 | 1 | 234.86 | TC,LDL | DETECTED | NM | B |
| *PIGV* | rs12748152 | 1 | 27.14 | HDL, LDL, TG | DETECTED | M | A |
| *APOB* | rs1367117 | 2 | 21.2600 | LDL,TC | DETECTED | M | B |
| *GCKR* | rs1260326 | 2 | 27.7300 | TG,TC | DETECTED | M | B |
| *IRS1* | rs2972146 | 2 | 227.1000 | HDL,TG | DETECTED | NA | B |
| *INSIG2* | rs10490626 | 2 | 118.84 | LDL, TCa | DETECTED | M | B |
| *ABCG5/8* | rs4299376 | 2 | 44.0700 | LDL,TC | DETECTED | M | B |
| *LOC84931* | rs2030746 | 2 | 121.31 | LDL, TC | DETECTED | NM | A |
| *UGT1A1* | rs11563251 | 2 | 234.68 | TC, LDL | DETECTED | M | A |
| *DNAJC13* | rs17404153 | 3 | 132.16 | LDL, HDLb | DETECTED | M | C |
| *CMTM6* | rs7640978 | 3 | 32.53 | LDL, TC | DETECTED | M | A |
| *LRPAP1* | rs6831256 | 4 | 3.47 | TG, TCa, LDLa | DETECTED | M | A |
| *HMGCR* | rs12916 | 5 | 74.6600 | TC,LDL | DETECTED | M | A |
| *TIMD4* | rs6882076 | 5 | 156.3900 | TC,TG,LDL | DETECTED | M | A |
| *CSNK1G3* | rs4530754 | 5 | 122.86 | LDL, TC | DETECTED | M | A |
| *LPA* | rs1564348 | 6 | 160.5800 | LDL,TC | DETECTED | M | B |
| *RSPO3* | rs1936800 | 6 | 127.44 | HDL, TGa | DETECTED | M | B |
| *HLA* | rs3177928 | 6 | 32.4100 | TC,LDL | DETECTED | M | B |
| *FRK* | rs9488822 | 6 | 116.3100 | TC,LDL | DETECTED | M | B |
| *MYLIP* | rs3757354 | 6 | 16.1300 | LDL,TC | DETECTED | M | A |
| *HFE* | rs1800562 | 6 | 26.0900 | LDL,TC | DETECTED | M | A |
| *VEGFA* | rs998584 | 6 | 43.76 | TG, HDL | DETECTED | M | A |
| *MIR148A* | rs4722551 | 7 | 25.99 | LDL, TGc, TC | DETECTED | M | B |
| *NPC1L1* | rs2072183 | 7 | 44.5800 | TC,LDL | DETECTED | M | B |
| *MLXIPL* | rs17145738 | 7 | 72.9800 | TG,HDL | DETECTED | M | B |
| *DNAH11* | rs12670798 | 7 | 21.6100 | TC,LDL | DETECTED | M | A |
| *PPP1R3B* | rs9987289 | 8 | 9.1800 | HDL,TC,LDL | DETECTED | NM | B |
| *LPL* | rs12678919 | 8 | 19.8400 | TG,HDL | DETECTED | M | B |
| *TRIB1* | rs2954029 | 8 | 126.4900 | TG,TC,LDL,HDL | DETECTED | M | B |
| *NAT2* | rs1495741 | 8 | 18.2700 | TG,TC | DETECTED | M | B |
| *CYP7A1* | rs2081687 | 8 | 59.3900 | TC,LDL | DETECTED | M | B |
| *PLEC1* | rs11136341 | 8 | 145.0400 | LDL,TC | DETECTED | M | B |
| *SOX17* | rs10102164 | 8 | 55.42 | LDL, TC | DETECTED | M | A |
| *TTC39B* | rs581080 | 9 | 15.3100 | HDL,TC | DETECTED | M | B |
| *ABO* | rs9411489 | 9 | 136.1550 | LDL,TC | DETECTED | M | B |
| *ABCA1* | rs1883025 | 9 | 107.6600 | HDL,TC | DETECTED | M | A |
| *VLDLR* | rs3780181 | 9 | 2.64 | TC, LDL | DETECTED | M | A |
| *GPAM* | rs2255141 | 10 | 113.9300 | TC,LDL | DETECTED | M | B |
| *8-Mar* | rs970548 | 10 | 46.01 | HDL, TC | DETECTED | M | B |
| *APOA1* | rs964184 | 11 | 116.6500 | TG,TC,HDL,LDL | DETECTED | M | B |
| *ST3GAL4* | rs11220462 | 11 | 126.2400 | LDL,TC | DETECTED | M | B |
| *FADS1-2-3* | rs174546 | 11 | 61.5700 | TG,LDL,TC,HDL | DETECTED | M | B |
| *UBASH3B* | rs7941030 | 11 | 122.5200 | TC,HDL | DETECTED | M | B |
| *LRP1* | rs11613352 | 12 | 57.7900 | TG,HDL | DETECTED | NM | B |
| *BRAP* | rs11065987 | 12 | 112.0700 | TC,LDL | DETECTED | M | B |
| *HNF1A* | rs1169288 | 12 | 121.4200 | TC,LDL | DETECTED | M | B |
| *ZNF664* | rs4765127 | 12 | 124.4600 | HDL,TG | DETECTED | M | B |
| *LIPC* | rs1532085 | 15 | 58.6800 | HDL,TC,TG | DETECTED | M | C (level 1) |
| *FTO* | rs1121980 | 16 | 53.81 | HDL, TGb | DETECTED | M | C (level 1) |
| *CETP* | rs3764261 | 16 | 56.9900 | HDL,LDL,TC,TG | DETECTED | M | B |
| *HPR* | rs2000999 | 16 | 72.1100 | TC,LDL | DETECTED | M | A |
| *OSBPL7* | rs7206971 | 17 | 45.4300 | LDL,TC | DETECTED | NM | B |
| *DLG4* | rs314253 | 17 | 7.09 | TC, LDL | DETECTED | M | A |
| *LIPG* | rs7241918 | 18 | 47.1600 | HDL,TC | DETECTED | M | B |
| *APOE* | rs4420638 | 19 | 45.4200 | LDL,TC,HDL | DETECTED | M | B |
| *LDLR* | rs6511720 | 19 | 11.2000 | LDL,TC | DETECTED | M | A |
| *CILP2* | rs10401969 | 19 | 19.4100 | TC,TG,LDL | DETECTED | NM | A |
| *PEPD* | rs731839 | 19 | 33.9 | TG, HDL | DETECTED | M | A |
| *MAFB* | rs2902940 | 20 | 39.0900 | TC,LDL | DETECTED | NM | C(level 5) |
| *TOP1* | rs6029526 | 20 | 39.6700 | LDL,TC | DETECTED | M | B |
| *PLTP* | rs6065906 | 20 | 44.5500 | HDL,TG | DETECTED | M | B |
| *HNF4A* | rs1800961 | 20 | 43.0400 | HDL,TC | DETECTED | M | A |
| *PPARA* | rs4253772 | 22 | 46.63 | TC, LDLa | DETECTED | M | B |

In this table, the columns in pheGWAS, gene match and SNP match corresponds to the PheGWAS results we obtained.

Match (M); if the gene symbol in a particular PheGWAS base pair interval segment is the same trait as reported in the GLGC
No Match (NM); if the gene is not identified by PheGWAS.

A - SNP identical with GLGC and PheGWAS
B – GLGC SNP in LD with SNP identified by PheGWAS
C – SNP identified by SNP thinning

**Supplementary Table 2b:** Summary of comparative analysis of loci identified by GLGC and PheGWAS results for association of loci with multiple traits

| | |
|---|---|
| Loci identified by GLGC | 69 |
| Total Loci Detected by PheGWAS | 69 |
| Gene Match | 59 |
| Gene No Match | 9 |
| Gene not mapped | 1 |
| SNP’s in Category A | 21 |
| SNP’s in Category B | 44 |
| SNP’s in Category C | 4 |

**Supplementary Table 2c:** LD Score between SNPs in GLGC and PheGWAS in multiple traits

| Chromosome | Position_MB | Multiple Traits | SNP Match | SNP ID | SNP ID phegwas | d' | r2 |
|---|---|---|---|---|---|---|---|
| 1 | 25.78 | TC,LDL | M | rs12027135 | rs11802413,rs10903129 | 1,1 | 0.98,0.98 |
| 1 | 55.50 | LDL,TC | NM | rs2479409 | rs11591147,rs11591147 | 1,1 | 0.01,0.01 |
| 1 | 63.03 | TG,LDL,TC | M | rs2131925 | rs4587594,rs11485618,rs3850634 | 1,1,1 | 0.99,0.98 |
| 1 | 109.82 | LDL,TC | M | rs629301 | rs646776,rs646776 | 1,1 | 0.99,0.99 |
| 1 | 220.97 | TC,LDL | M | rs2642442 | rs2642438,rs2642438 | 0.92,0.92 | 0.80,0.80 |
| 1 | 230.30 | HDL,TG | M | rs4846914 | rs1321257 | 0.99 | 0.95 |
| 1 | 234.86 | TC,LDL | M | rs514230 | rs558971,rs2587534 | 0.97,0.97 | 0.94,0.93 |
| 2 | 21.2600 | LDL,TC | PM | rs1367117 | rs515135 | 0.82 | 0.07 |
| 2 | 27.7300 | TG,TC | M | rs1260326 | rs780093,rs780093 | 0.95,0.95 | 0.91,0.91 |
| 2 | 227.1000 | HDL,TG | M | rs2972146 | rs1515110 | 0.94 | 0.88 |
| 2 | 118.84 | LDL, TCa | M | rs10490626 | rs17526895 | 1 | 0.99 |
| 2 | 44.0700 | LDL,TC | M | rs4299376 | rs6544713,rs6544713 | 0.99,0.99 | 0.97,0.97 |
| 3 | 132.16 | LDL, HDLb | PM | rs17404153 | NA | NA | NA |
| 6 | 160.5800 | LDL,TC | M | rs1564348 | rs11753995 | 0.98 | 0.95 |
| 6 | 127.44 | HDL, TGa | M | rs1936800 | rs719726 | 0.95 | 0.75 |
| 6 | 32.4100 | TC,LDL | PM | rs3177928 | rs9391858,rs10947332 | 0.96,0.89 | 0.85,0.56 |
| 6 | 116.3100 | TC,LDL | NM | rs9488822 | rs11153594,rs6909746 | 0.95,0.95 | 0.66,0.66 |
| 7 | 25.99 | LDL, TGc, TC | PM | rs4722551 | rs4719841 | 0.88 | 0.1 |
| 7 | 44.5800 | TC,LDL | NM | rs2072183 | rs2073547,rs2073547 | 0.96,0.96 | 0.73,0.73 |
| 7 | 72.9800 | TG,HDL | PM | rs17145738 | rs11974409 | 0.99 | 0.55 |
| 8 | 9.1800 | HDL,TC,LDL | M | rs9987289 | rs4240624 | 1 | 1 |
| 8 | 19.8400 | TG,HDL | PM | rs12678919 | rs13702 | 0.99 | 0.31 |
| 8 | 126.4900 | TG,TC,LDL,HDL | M | rs2954029 | rs2954022,rs10808546 | 1,0.98 | 0.98,0.88 |
| 8 | 18.2700 | TG,TC | PM | rs1495741 | rs4921914,rs1961456 | 1,0.94 | 1,0.70 |
| 8 | 59.3900 | TC,LDL | M | rs2081687 | rs4738684,rs13277801 | 0.97,0.97 | 0.93,0.94 |
| 8 | 145.0400 | LDL,TC | M | rs11136341 | rs7832643,rs7832643 | 0.92,0.92 | 0.78,0.78 |
| 9 | 15.3100 | HDL,TC | PM | rs581080 | rs686030 | 0.96 | 0.62 |
| 9 | 136.1550 | LDL,TC | M | rs9411489 | rs579459,rs579459 | 1,1 | 0.83,0.83 |
| 10 | 113.9300 | TC,LDL | M | rs2255141 | rs2419604 | 0.99 | 0.97 |
| 11 | 116.6500 | TG,TC,HDL,LDL | PM | rs964184 | rs10790162 | 0.99 | 0.51 |
| 11 | 126.2400 | LDL,TC | M | rs11220462 | rs10893499 | 1 | 0.99 |
| 11 | 61.5700 | TG,LDL,TC,HDL | M | rs174546 | rs174535,rs174583,rs1535,rs102275 | 1,0.99,0.99,1 | 0.97,0.92,0.97,0.93 |
| 11 | 122.5200 | TC,HDL | M | rs7941030 | rs7117842,rs7117842 | 0.99,0.99 | 0.95,0.95 |
| 12 | 57.7900 | TG,HDL | M | rs11613352 | rs3741414 | 1 | 0.99 |
| 12 | 112.0700 | TC,LDL | M | rs11065987 | rs653178,rs653178 | 0.97,0.97 | 0.78,0.78 |
| 12 | 121.4200 | TC,LDL | M | rs1169288 | rs2244608 | 0.99 | 0.96 |
| 12 | 124.4600 | HDL,TG | M | rs4765127 | rs7973683,rs11057408 | 0.99,1 | 0.97,1 |
| 15 | 58.6800 | HDL,TC,TG | NM | rs1532085 | rs10468017,rs10468017,rs588136 | 0.98,0.98,0.001 | 0.67,0.67,0 |
| 16 | 53.81 | HDL, TGb | PM | rs1121980 | rs9930333 | NA | NA |
| 16 | 56.9900 | HDL,LDL,TC,TG | PM | rs3764261 | rs9989419,rs247616,rs247616,rs1800775 | 0.81,0.99,0.99,1 | 0.18,0.97,0.97,0.43 |
| 17 | 45.4300 | LDL,TC | M | rs7206971 | rs6504872,rs6504872 | 1,1 | 1,1 |
| 18 | 47.1600 | HDL,TC | M | rs7241918 | rs4939883,rs2156552 | 0.99,0.98 | 0.92,0.94 |
| 19 | 45.4200 | LDL,TC,HDL | NM | rs4420638 | rs7254892,rs7412,rs2075650 | 0.84,0.92,0.72 | 0.01,0.01,0.31 |
| 20 | 39.0900 | TC,LDL | NM | rs2902940 | rs2235367,rs6065311 | 0.02,0.02 | 0.0001,0.0001 |
| 20 | 39.6700 | LDL,TC | PM | rs6029526 | rs2235367,rs6065312 | 0.99,1 | 0.96,0.22 |
| 20 | 44.5500 | HDL,TG | PM | rs6065906 | rs4465830,rs4810479 | 0.99,1 | 0.99,0.63 |
| 22 | 46.63 | TC, LDLa | M | rs4253772 | rs4253776 | 0.99 | 0.95 |

**Supplementary Table 3:** PheGWAS findings showcasing Chromosome Number, Marker Name, Associated Traits, Genes and the respective p-values for Chromosome 1

| Chr | Position group (Mb) | Marker Name | Associated Traits | Genes | P-Values | -log10 (P-value) |
|---|---|---|---|---|---|---|
| 1 | 23 | rs1077514 | TOTAL_CHOLESTROL | ASAP3 RP4-654C18.1 TCEA3 | 6.4e-09 | 8.19 |
| 1 | 25 | rs10903129 rs11802413 | LDL TOTAL_CHOLESTROL | AL031284.1 RHCE RP3-469D22.1 TMEM57 | 3.03e-17 1.576e-14 | 16.52 13.8 |
| 1 | 26 | rs2229714 | HDL | MIR1976 RN7SL679P RPS6KA1 Y_RNA | 6.094e-09 | 8.22 |
| 1 | 27 | rs12748152 rs12748152 rs12748152 | HDL LDL TRIGS | ARID1A PIGV RN7SL165P RN7SL501P ZDHHC18 | 9.737e-16 3.209e-12 1.1e-09 | 15.01 11.49 8.96 |
| 1 | 39 | rs2296173 rs16837533 | HDL TRIGS | BMP8A KIAA0754 MACF1 RP11-416A14.1 RP11-420K8.1 | 2.803e-17 3.873e-08 | 16.55 7.41 |
| 1 | 40 | rs4660293 rs17513135 | HDL TRIGS | BMP8A OXCT2P1 PABPC4 PPIEL RP11-69E11.4 RP11-69E11.8 SNORA55 | 2.863e-18 1.633e-08 | 17.54 7.79 |
| 1 | 55 | rs11591147 rs11591147 | LDL TOTAL_CHOLESTROL | BSND PCSK9 RP11-12C17.2 TMEM61 USP24 | 8.58e-143 8.827e-86 | 142.07 85.05 |
| 1 | 56 | rs12066643 | LDL | RP11-466L17.1 RP11-90C4.1 | 1.063e-08 | 7.97 |
| 1 | 62 | rs1168013 rs1168032 rs1168013 | LDL TRIGS TOTAL_CHOLESTROL | DOCK7 RP11-293K19.1 | 5.713e-33 1.486e-80 1.544e-80 | 32.24 79.83 79.81 |
| 1 | 63 | rs11485618 rs4587594 rs3850634 | LDL TRIGS TOTAL_CHOLESTROL | AL138847.1 ANGPTL3 DOCK7 RP5-849H19.2 RP11-230B22.1 | 3.729e-33 3.503e-82 1.39e-81 | 32.43 81.46 80.86 |
| 1 | 92 | rs4970712 rs6603981 | LDL TOTAL_CHOLESTROL | EVI5 GFI1 | 2.458e-13 7.846e-15 | 12.61 14.11 |
| 1 | 93 | rs12133576 rs1556562 rs1556562 | HDL LDL TOTAL_CHOLESTROL | DR1 RP4-713B5.2 RP4-717I23.3 Y_RNA EVI5 RP4-593M8.1 | 6.15e-11 4.558e-13 1.398e-14 | 10.21 12.34 13.85 |
| 1 | 109 | rs12740374 rs646776 rs646776 | HDL LDL TOTAL_CHOLESTROL | CELSR2 MYBPHL PSRC1 SARS SORT1 | 1.687e-15 1.63e-272 4.77e-187 | 14.77 271.79 186.32 |
| 1 | 110 | rs333947 rs650985 rs518076 | HDL LDL TOTAL_CHOLESTROL | CSF1 RP11-195M16.1 RP11-195M16.3 AC000032.2 AMPD2 GNAT2 GSTM1 GSTM2 GSTM4 RP5-1160K1.1 RP5-1160K1.6 GNAI3 GPR61 MIR197 RNU6V RP5-1160K1.3 RP5-1160K1.8 | 3.166e-09 1.899e-15 5.058e-10 | 8.5 14.72 9.3 |
| 1 | 150 | rs267733 | LDL | ANXA9 CERS2 FAM63A PRUNE RNU6-884P RP11-316M1.12 RP11-316M1.3 SETDB1 | 5.285e-09 | 8.28 |
| 1 | 156 | rs12145743 | HDL | CRABP2 HDGF ISG20L2 MRPL24 PRCC RP11-66D17.3 RP11-66D17.5 RRNAD1 | 1.803e-08 | 7.74 |
| 1 | 178 | rs4650994 | HDL | C1ORF220 C1orf220 LINC00083 RNA5SP69 TEX35 | 6.696e-09 | 8.17 |
| 1 | 182 | rs1689797 | HDL | GS1-122H1.2 GS1-122H1.3 | 2.852e-21 | 20.54 |
| 1 | 220 | rs2642438 rs2642438 rs2642438 | HDL LDL TOTAL_CHOLESTROL | HLA-AS1 MARC1 MARC2 RNU6ATAC35P RP11-295M18.2 RP11-295M18.6 | 7.781e-14 7.318e-16 1.283e-18 | 13.11 15.14 17.89 |
| 1 | 221 | rs2247213 rs11590084 | LDL TOTAL_CHOLESTROL | HLA-AS1 HLX RP11-295M18.2 RNU6ATAC35P RP11-295M18.6 | 1.412e-08 8.387e-08 | 7.85 7.08 |
| 1 | 230 | rs4846914 rs1321257 | HDL TRIGS | GALNT2 | 3.513e-41 5.986e-31 | 40.45 30.22 |
| 1 | 234 | rs2587534 rs558971 | LDL TOTAL_CHOLESTROL | RP4-781K5.4 RP4-781K5.5 RP4-781K5.6 RP4-781K5.7 RP4-781K5.8 | 8.055e-25 7.025e-28 | 24.09 27.15 |

**Supplementary Table 4:** PheGWAS findings showcasing Chromosome Number, Marker Name, Associated Traits, Genes and the respective p-values for Chromosome 2

| Chr | Position group (Mb) | Marker Name | Associated Traits | Genes | P-Values | -log10 (P-value) |
|---|---|---|---|---|---|---|
| 2 | 20 | rs11679386 rs11679386 | LDL TOTAL_CHOLESTROL | C2orf43 | 4.746e-14 2.226e-11 | 13.32 10.65 |
| 2 | 21 | rs676210 rs1367117 rs676210 rs515135 | HDL LDL TRIGS TOTAL_CHOLESTROL | APOB RP11-116D2.1 | 2.345e-54 9.48e-183 3.284e-71 6.38e-151 | 53.63 182.02 70.48 150.2 |
| 2 | 27 | rs780093 rs1260326 rs780093 | LDL TRIGS TOTAL_CHOLESTROL | AC109829.1 FNDC4 GCKR IFT172 RNU6-986P | 2.359e-08 2.29e-239 2.591e-42 | 7.63 238.64 41.59 |
| 2 | 28 | rs13030345 rs13030345 | TRIGS TOTAL_CHOLESTROL | AC110084.1 MRPL33 RBKS | 1.019e-57 2.787e-10 | 56.99 9.55 |
| 2 | 43 | rs10495907 rs10495907 | LDL TOTAL_CHOLESTROL | ABCG5 DYNC2LI1 PLEKHH2 RN7SKP66 | 1.264e-13 3.085e-11 | 12.9 10.51 |
| 2 | 44 | rs6544713 rs6544713 | LDL TOTAL_CHOLESTROL | ABCG5 ABCG8 DYNC2LI1 LRPPRC RNU6-1048P | 4.843e-83 1.685e-81 | 82.31 80.77 |
| 2 | 62 | rs1534420 | LDL | AC092155.4 EHBP1 | 3.896e-08 | 7.41 |
| 2 | 63 | rs2710642 | LDL | EHBP1 RP11-443F16.1 | 6.089e-09 | 8.22 |
| 2 | 118 | rs10490626 rs17526895 | LDL TOTAL_CHOLESTROL | INSIG2 AC009303.1 CCDC93 RN7SL111P | 1.697e-12 5.777e-09 | 11.77 8.24 |
| 2 | 121 | rs2030746 rs2030746 | LDL TOTAL_CHOLESTROL | AC073257.1 AC073257.2 | 8.605e-09 3.603e-08 | 8.07 7.44 |
| 2 | 135 | rs16831243 rs6730157 | LDL TOTAL_CHOLESTROL | CCNT2 MAP3K19 RAB3GAP1 SNORA40 ZRANB3 | 9.063e-12 1.179e-13 | 11.04 12.93 |
| 2 | 136 | rs4988235 rs4988235 | LDL TOTAL_CHOLESTROL | AC011893.3 AC011999.1 LCT MCM6 Y_RNA | 3.218e-11 3.975e-14 | 10.49 13.4 |
| 2 | 165 | rs7607980 rs10195252 rs13389219 | HDL LDL TRIGS | AC019181.3 COBLL1 SNORA70F GRB14 | 1.807e-15 3.812e-08 2.598e-15 | 14.74 7.42 14.59 |
| 2 | 169 | rs2287623 rs2287623 | LDL TOTAL_CHOLESTROL | ABCB11 | 5.399e-08 4.085e-12 | 7.27 11.39 |
| 2 | 203 | rs934287 rs11694172 | LDL TOTAL_CHOLESTROL | ICA1L KRT8P15 WDR12 AC009960.3 AC009960.6 AC009960.7 FAM117B MTND4P30 | 1.214e-07 1.951e-09 | 6.92 8.71 |
| 2 | 204 | rs1174604 | TOTAL_CHOLESTROL | RAPH1 | 3.115e-07 | 6.51 |
| 2 | 211 | rs1047891 | HDL | CPS1 | 8.73e-10 | 9.06 |
| 2 | 216 | rs1250229 rs1250229 | LDL TOTAL_CHOLESTROL | AC012462.1 AC012462.2 AC012462.3 FN1 | 3.13e-08 2.382e-07 | 7.5 6.62 |
| 2 | 227 | rs1515110 rs2972146 | HDL TRIGS | | 8.038e-18 2.974e-15 | 17.09 14.53 |
| 2 | 234 | rs11563251 rs11563251 | LDL TOTAL_CHOLESTROL | AC114812.10 AC114812.5 AC114812.9 DNAJB3 MROH2A RPL17P11 UGT1A1 UGT1A10 UGT1A2P UGT1A3 UGT1A4 UGT1A5 UGT1A6 UGT1A7 UGT1A8 UGT1A9 | 4.499e-08 1.266e-09 | 7.35 8.9 |

**Supplementary Table 5:** PheGWAS findings showcasing Chromosome Number, Marker Name, Associated Traits, Genes and the respective p-values for Chromosome 3

| Chr | Position group (Mb) | Marker Name | Associated Traits | Genes | P-Values | -log10 (P-value) |
|---|---|---|---|---|---|---|
| 3 | 11 | rs2606736 | HDL | ATG7 | 4.799e-08 | 7.32 |
| 3 | 12 | rs2292101 rs9875338 rs10440120 rs7616006 | HDL LDL TRIGS TOTAL_CHOLESTROL | PPARG GSTM5P1 TSEN2 SYN2 | 1.359e-07 2.21e-11 5.343e-11 8.406e-17 | 6.87 10.66 10.27 16.08 |
| 3 | 32 | rs7640978 rs7640978 | LDL TOTAL_CHOLESTROL | AC104306.1 AC104306.2 CMTM6 CMTM7 DYNC1LI1 | 9.837e-09 1.658e-08 | 8.01 7.78 |
| 3 | 47 | rs2290547 | HDL | CCDC12 NBEAL2 NRADDP SETD2 | 3.69e-09 | 8.43 |
| 3 | 49 | rs7613875 | HDL | CTD-2330K9.2 CTD-2330K9.3 MON1A MST1R RBM6 | 1.792e-11 | 10.75 |
| 3 | 50 | rs2013208 | HDL | RBM5 RBM6 RP11-493K19.3 | 8.916e-12 | 11.05 |
| 3 | 52 | rs13326165 | HDL | NISCH NT5DC2 PBRM1 SMIM4 STAB1 TNNC1 | 9.042e-11 | 10.04 |
| 3 | 53 | rs2336725 | HDL | AC096887.1 RFT1 RP11-894J14.5 SERBP1P3 SFMBT1 | 2.316e-07 | 6.64 |
| 3 | 58 | rs13315871 | TOTAL_CHOLESTROL | PDHB PXK RP11-802O23.3 | 3.48e-08 | 7.46 |
| 3 | 119 | rs6805251 | HDL | GSK3B NR1I2 PHBP8 | 1.332e-08 | 7.88 |
| 3 | 131 | rs13076253 | HDL | CPNE4 MIR5704 | 4.961e-09 | 8.3 |
| 3 | 132 | rs17404153 rs17404153 | LDL TOTAL_CHOLESTROL | DNAJC13 NIP7P2 | 1.832e-09 1.362e-07 | 8.74 6.87 |
| 3 | 135 | rs687339 rs645040 | HDL TRIGS | MSL2 PCCB RP11-463H24.1 | 7.108e-13 1.83e-12 | 12.15 11.74 |
| 3 | 136 | rs1279840 rs12630999 | HDL TRIGS | PCCB STAG1 HMGN1P10 RNY4P4 | 2.135e-12 3.151e-10 | 11.67 9.5 |
| 3 | 142 | rs4683438 | TOTAL_CHOLESTROL | PAQR9 PCOLCE2 RP11-372E1.4 RP11-372E1.6 RP11-372E1.7 U2SURP | 2.509e-07 | 6.6 |
| 3 | 156 | rs1482852 | HDL | LEKR1 LINC00880 LINC00881 RP11-6F2.5 | 6.34e-08 | 7.2 |
| 3 | 170 | rs10513688 | TRIGS | RNU1-70P SLC2A2 | 1.543e-07 | 6.81 |

**Supplementary Table 6:** PheGWAS findings showcasing Chromosome Number, Marker Name, Associated Traits, Genes and the respective p-values for Chromosome 4

| Chr | Position group (Mb) | Marker Name | Associated Traits | Genes | P-Values | -log10 (P-value) |
|---|---|---|---|---|---|---|
| 4 | 3 | rs6818397 rs6831256 rs6818397 | LDL TRIGS TOTAL_CHOLESTROL | DOK7 HGFAC RGS12 AL590235.1 LRPAP1 RP11-529E10.6 | 1.677e-08 1.602e-12 9.51e-11 | 7.78 11.8 10.02 |
| 4 | 26 | rs10019888 | HDL |  | 4.901e-08 | 7.31 |
| 4 | 69 | rs176813 | TOTAL_CHOLESTROL | RP11-1267H10.4 | 1.178e-07 | 6.93 |
| 4 | 87 | rs3775228 rs3736669 | HDL TRIGS | AFF1 | 4.53e-09 6.542e-15 | 8.34 14.18 |
| 4 | 88 | rs442177 rs442177 | HDL TRIGS | AFF1 | 2.193e-09 1.316e-18 | 8.66 17.88 |
| 4 | 89 | rs3822072 rs13133548 | HDL TRIGS | FAM13A | 4.058e-12 2.108e-07 | 11.39 6.68 |
| 4 | 100 | rs2602836 | HDL | ADH4 ADH5 METAP1 RP11-571L19.8 RP11-696N14.1 | 4.964e-08 | 7.3 |
| 4 | 102 | rs17199964 | HDL | BANK1 | 1.843e-07 | 6.73 |
| 4 | 103 | rs13107325 | HDL | SLC39A8 | 1.065e-15 | 14.97 |
| 4 | 157 | rs6822892 | HDL | PDGFC RP11-154F14.2 | 1.933e-07 | 6.71 |

**Supplementary Table 7:** PheGWAS findings showcasing Chromosome Number, Marker Name, Associated Traits, Genes and the respective p-values for Chromosome 5

| Chr | Position group (Mb) | Marker Name | Associated Traits | Genes | P-Values | -log10 (P-value) |
|---|---|---|---|---|---|---|
| 5 | 53 | rs6450176 | HDL | ARL15 RN7SL801P | 6.875e-10 | 9.16 |
| 5 | 55 | rs3936511 rs9686661 | HDL TRIGS | AC022431.2 AC022431.3 | 2.959e-09 2.541e-16 | 8.53 15.59 |
| 5 | 67 | rs4976033 | HDL | CTC-537E7.3 CTD-2582M21.1 | 6.421e-08 | 7.19 |
| 5 | 74 | rs12916 rs12916 | LDL TOTAL_CHOLESTROL | COL4A3BP CTD-2235C13.1 CTD-2235C13.2 CTD-2235C13.3 HMGCR | 7.792e-78 4.547e-74 | 77.11 73.34 |
| 5 | 75 | rs2112347 rs2112347 | LDL TOTAL_CHOLESTROL | ANKDD1B POC5 RNU6-680P SLC25A5P9 | 4.43e-30 1.72e-33 | 29.35 32.76 |
| 5 | 122 | rs4530754 rs4530754 | LDL TOTAL_CHOLESTROL | CSNK1G3 Y_RNA | 3.576e-12 1.678e-09 | 11.45 8.78 |
| 5 | 156 | rs6882076 rs6882076 rs6882076 | LDL TRIGS TOTAL_CHOLESTROL | APOOP1 TIMD4 | 3.31e-31 1.513e-15 5.354e-41 | 30.48 14.82 40.27 |
| 5 | 173 | rs72812840 | TRIGS | C5orf47 CPEB4 | 2.896e-07 | 6.54 |

**Supplementary Table 8:** PheGWAS findings showcasing Chromosome Number, Marker Name, Associated Traits, Genes and the respective p-values for Chromosome 6

| Chr | Position group (Mb) | Marker Name | Associated Traits | Genes | P-Values | -log10 (P-value) |
|---|---|---|---|---|---|---|
| 6 | 16 | rs3757354 rs3757354 | LDL TOTAL_CHOLESTROL | MIR4639 MRPL42P2 MYLIP RP1-13D10.2 RP1-13D10.3 RP1-13D10.5 U3 | 2.087e-17 2.215e-15 | 16.68 14.65 |
| 6 | 25 | rs17526722 rs1408272 rs1408272 | HDL LDL TOTAL_CHOLESTROL | HIST1H2APS2 SLC17A2 SLC17A3 TRIM38 SLC17A1 | 1.42e-07 3.675e-09 5.034e-08 | 6.85 8.43 7.3 |
| 6 | 26 | rs1800562 rs1800562 | LDL TOTAL_CHOLESTROL | HFE HIST1H1C HIST1H1T HIST1H2AC HIST1H2BB HIST1H2BC HIST1H3C HIST1H4C U91328.2 | 8.253e-14 1.905e-12 | 13.08 11.72 |
| 6 | 29 | rs5013093 | TRIGS | HCG4P7 HCG4P8 HLA-G HLA-H HLA-T MICF MICG RPL7AP7 | 1.358e-07 | 6.87 |
| 6 | 30 | rs1634721 rs1634721 | TRIGS TOTAL_CHOLESTROL | HCG22 MUC21 MUC22 | 4.015e-13 5.279e-12 | 12.4 11.28 |
| 6 | 31 | rs2844513 rs10947207 rs2247056 rs2247056 | HDL LDL TRIGS TOTAL_CHOLESTROL | AL671883.1 FGFR3P1 HCP5 HLA-S MICA XXbac-BPG181B23.4 XXbac-BPG181B23.6 XXbac-BPG181B23.7 Y_RNA ZDHHC20P2 DHFRP2 HLA-B RNU6-283P XXbac-BPG248L24.12 HLA-C RPL3P2 USP8P1 WASF5P XXbac-BPG248L24.10 XXbac-BPG248L24.13 | 2.465e-09 2.448e-12 3.861e-21 2.203e-20 | 8.61 11.61 20.41 19.66 |
| 6 | 32 | rs1980493 rs10947332 rs11752643 rs9391858 | HDL LDL TRIGS TOTAL_CHOLESTROL | BTNL2 C6orf10 HCG23 HLA-DRA RNU6-603P HLA-DQA2 HLA-DQB1 HLA-DQB1-AS1 HLA-DQB2 HLA-DQB3 MIR3135B MTCO3P1 XXbac-BPG254F23.6 XXbac-BPG254F23.7 HNRNPA1P2 | 3.764e-10 6.969e-18 3.96e-19 7.204e-22 | 9.42 17.16 18.4 21.14 |
| 6 | 34 | rs205262 rs3800457 rs2814982 | HDL LDL TOTAL_CHOLESTROL | C6orf106 RP3-391O22.2 RP3-391O22.3 SPDEF RN7SL200P RP11-140K17.2 RP11-140K17.3 RP3-375P9.2 RP3-391O22.1 snoU13 PACSIN1 | 3.878e-13 2.3e-07 3.678e-15 | 12.41 6.64 14.43 |
| 6 | 35 | rs3822921 rs3800406 rs3800406 | HDL LDL TOTAL_CHOLESTROL | AL138721.1 ANKS1A TCP11 SCUBE3 | 2.349e-09 9.392e-08 1.014e-13 | 8.63 7.03 12.99 |
| 6 | 39 | rs2758886 rs2758886 | LDL TOTAL_CHOLESTROL | KCNK16 KCNK17 KIF6 | 1.845e-07 3.008e-08 | 6.73 7.52 |
| 6 | 43 | rs998584 rs998584 | HDL TRIGS | RP1-261G23.7 VEGFA | 2.269e-11 3.424e-15 | 10.64 14.47 |
| 6 | 109 | rs884366 | HDL | C6orf183 CCDC162P PTCHD3P3 RP11-425D10.1 snoU13 | 1.671e-07 | 6.78 |
| 6 | 116 | rs6909746 rs11153594 | LDL TOTAL_CHOLESTROL | FRK TPI1P3 | 7.86e-11 1.266e-14 | 10.1 13.9 |
| 6 | 127 | rs1936800 rs719726 | HDL TRIGS | RSPO3 | 3.055e-10 2.486e-08 | 9.51 7.6 |
| 6 | 135 | rs9376090 | TOTAL_CHOLESTROL | CTA-212D2.2 HBS1L | 2.595e-09 | 8.59 |
| 6 | 139 | rs3861397 rs634869 | HDL TRIGS | RP11-12A2.3 | 8.401e-11 1.776e-14 | 10.08 13.75 |
| 6 | 152 | rs12525163 | HDL | ESR1 | 1.524e-07 | 6.82 |
| 6 | 160 | rs9457931 rs1564348 rs2665357 rs11753995 | HDL LDL TRIGS TOTAL_CHOLESTROL | LPA LPAL2 IGF2R RP11-288H12.4 RP11-317M22.1 SLC22A1 SLC22A2 AL591069.1 SLC22A3 | 7.297e-13 2.762e-21 8.327e-10 1.839e-23 | 12.14 20.56 9.08 22.74 |
| 6 | 161 | rs1084651 rs10455872 rs10455872 | HDL LDL TOTAL_CHOLESTROL | LPA PLG RP1-81D8.3 RP1-81D8.4 | 1.347e-11 1.936e-15 7.236e-17 | 10.87 14.71 16.14 |

**Supplementary Table 9:** PheGWAS findings showcasing Chromosome Number, Marker Name, Associated Traits, Genes and the respective p-values for Chromosome 7

| Chr | Position group (Mb) | Marker Name | Associated Traits | Genes | P-Values | -log10 (P-value) |
|---|---|---|---|---|---|---|
| 7 | 1 | rs1997243 rs1997243 | HDL TOTAL_CHOLESTROL | AC073957.15 AC091729.7 AC091729.8 C7orf50 GPER1 GPR146 MIR339 RP11-449P15.1 | 2.27e-07 2.719e-10 | 6.64 9.57 |
| 7 | 6 | rs702485 | HDL | DAGLB KDELR2 RAC1 | 6.45e-12 | 11.19 |
| 7 | 16 | rs17286602 | HDL | AC006035.1 ISPD RP11-196O16.1 | 2.932e-07 | 6.53 |
| 7 | 17 | rs4142995 | HDL | SNX13 | 9.365e-12 | 11.03 |
| 7 | 21 | rs12670798 rs12670798 | LDL TOTAL_CHOLESTROL | DNAH11 | 4.812e-14 9.478e-17 | 13.32 16.02 |
| 7 | 25 | rs4722551 rs4719841 rs4722551 | LDL TRIGS TOTAL_CHOLESTROL | CTD-2227E11.1 MIR148A | 3.949e-14 8.864e-11 7.019e-09 | 13.4 10.05 8.15 |
| 7 | 44 | rs2073547 rs2073547 | LDL TOTAL_CHOLESTROL | AC004938.5 DDX56 NPC1L1 RNU6-1097P TMED4 | 1.923e-21 3.83e-21 | 20.72 20.42 |
| 7 | 50 | rs4917014 | HDL | AC020743.4 IKZF1 | 1.026e-08 | 7.99 |
| 7 | 71 | rs2909969 | TRIGS | TYW1B | 1.528e-07 | 6.82 |
| 7 | 72 | rs17145738 rs11974409 | HDL TRIGS | BAZ1B BCL7B MLXIPL TBL2 | 4.952e-13 1.36e-100 | 12.31 99.87 |
| 7 | 73 | rs17145750 rs17145750 | HDL TRIGS | MLXIPL TBL2 | 5.654e-12 1.03e-99 | 11.25 98.99 |
| 7 | 116 | rs38855 | TRIGS | MET | 2.109e-08 | 7.68 |
| 7 | 130 | rs11765979 rs287621 | HDL TRIGS | KLF14 RP11-36B6.1 | 3.111e-17 7.671e-09 | 16.51 8.12 |
| 7 | 150 | rs17173637 | HDL | AOC1 RP5-1051J4.4 TMEM176A TMEM176B | 1.899e-08 | 7.72 |

**Supplementary Table 10:** PheGWAS findings showcasing Chromosome Number, Marker Name, Associated Traits, Genes and the respective p-values for Chromosome 8

| Chr | Position group (Mb) | Marker Name | Associated Traits | Genes | P-Values | -log10 (P-value) |
|---|---|---|---|---|---|---|
| 8 | 8 | rs2976940 | TRIGS | CTA-398F10.1 CTA-398F10.2 SGK223 | 1.138e-08 | 7.94 |
| 8 | 9 | rs4240624 rs9987289 rs9693857 rs9987289 | HDL LDL TRIGS TOTAL_CHOLESTROL | RNU6-1151P RNU6-526P RP11-115J16.1 RP11-115J16.2 RP11-115J16.3 RP11-375N15.1 | 1.323e-45 8.529e-24 1.691e-08 1.844e-36 | 44.88 23.07 7.77 35.73 |
| 8 | 10 | rs7014168 rs6995541 | HDL TRIGS | CTD-2135J3.3 PINX1 SNORD112 SOX7 RP11-177H2.2 | 9.203e-10 1.343e-12 | 9.04 11.87 |
| 8 | 11 | rs1062219 | TRIGS | C8orf49 FDFT1 GATA4 NEIL2 RP11-297N6.4 SUB1P1 | 1.686e-09 | 8.77 |
| 8 | 18 | rs1961456 rs4921914 rs1961456 | LDL TRIGS TOTAL_CHOLESTROL | NAT2 NATP RP11-685B14.1 RP11-685B14.3 | 6.883e-08 4.867e-17 3.458e-16 | 7.16 16.31 15.46 |
| 8 | 19 | rs13702 rs12678919 | HDL TRIGS | LPL | 1.28e-160 1.82e-199 | 159.89 198.74 |
| 8 | 55 | rs10102164 rs10102164 | LDL TOTAL_CHOLESTROL | RP11-53M11.2 RP11-53M11.3 RP11-53M11.4 SOX17 | 3.742e-11 4.596e-11 | 10.43 10.34 |
| 8 | 59 | rs13277801 rs4738684 rs4738684 | LDL TRIGS TOTAL_CHOLESTROL | CYP7A1 RP11-114M5.1 UBXN2B RP11-114M5.3 | 3.994e-17 8.819e-09 1.116e-23 | 16.4 8.05 22.95 |
| 8 | 116 | rs2293889 rs2737252 rs2737252 | HDL LDL TOTAL_CHOLESTROL | TRPS1 | 4.271e-17 7.039e-14 1.634e-16 | 16.37 13.15 15.79 |
| 8 | 121 | rs4871137 | HDL | RP11-713M15.2 SNTB1 | 1.926e-07 | 6.72 |
| 8 | 126 | rs10808546 rs2954029 rs2954022 rs2954029 | HDL LDL TRIGS TOTAL_CHOLESTROL | RN7SL590P RP11-136O12.2 TRIB1 | 4.106e-30 2.095e-50 2.23e-113 2.421e-65 | 29.39 49.68 112.65 64.62 |
| 8 | 144 | rs10087900 rs6984820 rs6984820 | HDL LDL TOTAL_CHOLESTROL | GLI4 GPIHBP1 ZFP41 EPPK1 MIR661 PLEC | 2.174e-09 3.769e-09 1.634e-07 | 8.66 8.42 6.79 |
| 8 | 145 | rs7832643 rs7832643 | LDL TOTAL_CHOLESTROL | GRINA MIR661 PARP10 PLEC | 2.671e-17 3.121e-13 | 16.57 12.51 |

**Supplementary Table 11:** PheGWAS findings showcasing Chromosome Number, Marker Name, Associated Traits, Genes and the respective p-values for Chromosome 9

| Chr | Position group (Mb) | Marker Name | Associated Traits | Genes | P-Values | -log10 (P-value) |
|---|---|---|---|---|---|---|
| 9 | 2 | rs3780181 rs3780181 | LDL TOTAL_CHOLESTROL | RP11-125B21.2 VLDLR | 1.764e-09 6.668e-10 | 8.75 9.18 |
| 9 | 15 | rs686030 rs686030 rs581080 | HDL TRIGS TOTAL_CHOLESTROL | RN7SL157P TTC39B | 4.292e-27 2.226e-07 1.022e-13 | 26.37 6.65 12.99 |
| 9 | 16 | rs3927680 | LDL | BNC2 | 2.15e-07 | 6.67 |
| 9 | 19 | rs10757056 | TOTAL_CHOLESTROL | ACER2 DENND4C NDUFA5P3 RP11-363E7.4 RP11-513M16.7 RP11-513M16.8 RPS6 | 5.361e-08 | 7.27 |
| 9 | 107 | rs1883025 rs1883025 rs1883025 rs1883025 | HDL LDL TRIGS TOTAL_CHOLESTROL | ABCA1 RP11-217B7.2 | 1.496e-65 6.141e-11 2.908e-07 5.749e-53 | 64.83 10.21 6.54 52.24 |
| 9 | 136 | rs579459 rs579459 | LDL TOTAL_CHOLESTROL | ABO LCN1P2 RP11-430N14.4 SURF6 Y_RNA | 2.419e-44 8.828e-42 | 43.62 41.05 |

**Supplementary Table 12:** PheGWAS findings showcasing Chromosome Number, Marker Name, Associated Traits, Genes and the respective p-values for Chromosome 10

| Chr | Position group (Mb) | Marker Name | Associated Traits | Genes | P-Values | -log10 (P-value) |
|---|---|---|---|---|---|---|
| 10 | 5 | rs1832007 | TRIGS | AKR1C4 AKR1CL1 RP11-445P17.3 RP11-445P17.6 U8 | 1.718e-12 | 11.76 |
| 10 | 17 | rs10904908 rs10904908 | LDL TOTAL_CHOLESTROL | RP11-124N14.3 TRDMT1 VIM VIM-AS1 | 3.135e-07 2.601e-11 | 6.5 10.58 |
| 10 | 45 | rs10900221 rs10900221 | HDL TOTAL_CHOLESTROL | ALOX5 MARCH8 RP11-67C2.2 | 1.004e-09 7.963e-09 | 9 8.1 |
| 10 | 46 | rs970548 rs970548 | HDL TOTAL_CHOLESTROL | MARCH8 | 1.706e-10 8.431e-09 | 9.77 8.07 |
| 10 | 64 | rs7073746 rs7073746 | HDL TRIGS | JMJD1C NRBF2 RNU6-543P RP11-144G16.1 | 1.791e-08 1.234e-15 | 7.75 14.91 |
| 10 | 65 | rs10761771 rs10761762 | HDL TRIGS | JMJD1C JMJD1C-AS1 PRELID1P3 RP11-351O1.3 | 4.12e-09 1.061e-17 | 8.39 16.97 |
| 10 | 94 | rs8211 rs2068888 | HDL TRIGS | CYP26A1 CYP26C1 EXOC6 NIP7P1 RP11-348J12.2 | 8.049e-08 1.682e-11 | 7.09 10.77 |
| 10 | 113 | rs2250802 rs2419604 rs2250802 rs2255141 | HDL LDL TRIGS TOTAL_CHOLESTROL | GPAM | 2.022e-17 7.49e-14 1.21e-10 6.514e-16 | 16.69 13.13 9.92 15.19 |
| 10 | 114 | rs2148489 rs2148489 | HDL TOTAL_CHOLESTROL | GUCY2GP TECTB | 1.412e-10 2.366e-10 | 9.85 9.63 |
| 10 | 115 | rs7076938 | HDL | ADRB1 | 1.712e-07 | 6.77 |

**Supplementary Table 13:** PheGWAS findings showcasing Chromosome Number, Marker Name, Associated Traits, Genes and the respective p-values for Chromosome 11

| Chr | Position group (Mb) | Marker Name | Associated Traits | Genes | P-Values | -log10 (P-value) |
|---|---|---|---|---|---|---|
| 11 | 10 | rs2923084 | HDL | AMPD3 RNU6ATAC33P | 5.02e-08 | 7.3 |
| 11 | 14 | rs2303975 | HDL | RP11-21L19.1 RRAS2 SPON1 | 1.585e-07 | 6.8 |
| 11 | 18 | rs10832962 rs10832962 | LDL TOTAL_CHOLESTROL | RP11-137N23.1 RP11-504G3.4 SPTY2D1 SPTY2D1-AS1 UEVLD | 6.619e-14 1.539e-14 | 13.18 13.81 |
| 11 | 45 | rs74458891 | HDL | CTD-2210P24.1 CTD-2210P24.2 CTD-2210P24.3 | 6.492e-09 | 8.19 |
| 11 | 46 | rs3136441 rs6485690 | HDL TRIGS | ARHGAP1 ATG13 CKAP5 F2 MIR5582 SNORD67 ZNF408 | 6.759e-29 2.061e-07 | 28.17 6.69 |
| 11 | 47 | rs3847502 rs10501321 rs1681630 | HDL TRIGS TOTAL_CHOLESTROL | MADD MYBPC3 NR1H3 RP11-17G12.3 SPI1 ACP2 DDB2 PTPRJ RP11-793I11.1 | 3.306e-38 1.412e-08 6.929e-08 | 37.48 7.85 7.16 |
| 11 | 48 | rs7946766 rs4752805 | HDL TOTAL_CHOLESTROL | AC103828.1 PTPRJ | 4.967e-24 1.617e-09 | 23.3 8.79 |
| 11 | 49 | rs11040329 | HDL | CTD-2026G22.1 | 5.46e-12 | 11.26 |
| 11 | 50 | rs2201637 | HDL | RP11-227P3.1 RP11-347H15.1 | 5.273e-10 | 9.28 |
| 11 | 51 | rs11246602 | HDL | OR4C46 OR4C50P OR4C7P OR4R2P | 1.681e-10 | 9.77 |
| 11 | 55 | rs11229606 | HDL | OR4A10P OR4A13P OR4A14P OR4A17P OR4A50P OR4A9P OR4X7P RP11-131J4.12 | 3.117e-10 | 9.51 |
| 11 | 56 | rs6591323 | HDL | OR5AL1 OR5AL2P OR5M4P OR5M9 OR5R1 OR8L1P OR8U1 | 1.508e-08 | 7.82 |
| 11 | 61 | rs102275 rs174583 rs174535 rs1535 | HDL LDL TRIGS TOTAL_CHOLESTROL | DAGLA FADS1 FADS2 FEN1 MIR1908 MIR611 MYRF RP11-467L20.10 TMEM258 FADS3 | 6.404e-28 7.003e-41 1.733e-41 8.624e-39 | 27.19 40.15 40.76 38.06 |
| 11 | 65 | rs12801636 | HDL | AP001362.1 EHBP1L1 FAM89B KCNK7 MAP3K11 MIR4489 MIR4690 PCNXL3 RELA SIPA1 SSSCA1 | 3.147e-08 | 7.5 |
| 11 | 75 | rs499974 | HDL | CTD-2530H12.1 CTD-2530H12.2 DGAT2 MOGAT2 RN7SL786P | 1.12e-08 | 7.95 |
| 11 | 116 | rs964184 rs964184 rs10790162 rs964184 | HDL LDL TRIGS TOTAL_CHOLESTROL | AP006216.10 AP006216.11 AP006216.5 APOA4 APOA5 BUD13 ZNF259 | 6.086e-48 2.008e-26 1.1e-249 2.838e-55 | 47.22 25.7 248.96 54.55 |
| 11 | 117 | rs508487 rs508487 rs508487 rs508487 | HDL LDL TRIGS TOTAL_CHOLESTROL | PAFAH1B2 PCSK7 RNF214 SIDT2 TAGLN | 5.45e-11 1.773e-10 9.22e-119 1.002e-27 | 10.26 9.75 118.04 27 |
| 11 | 118 | rs11603023 | TOTAL_CHOLESTROL | AP002954.3 ARCN1 IFT46 PHLDB1 RNU6-1157P TREH | 1.151e-08 | 7.94 |
| 11 | 122 | rs7117842 rs7117842 | HDL TOTAL_CHOLESTROL | GLULP3 RP11-266E8.2 UBASH3B | 1.057e-14 2.479e-15 | 13.98 14.61 |
| 11 | 126 | rs17135399 rs10893499 rs11220462 | HDL LDL TOTAL_CHOLESTROL | DCPS RP11-712L6.5 ST3GAL4 TIRAP KIRREL3 | 4.26e-09 3.861e-21 5.487e-15 | 8.37 20.41 14.26 |

**Supplementary Table 14:** PheGWAS findings showcasing Chromosome Number, Marker Name, Associated Traits, Genes and the respective p-values for Chromosome 12

| Chr | Position group (Mb) | Marker Name | Associated Traits | Genes | P-Values | -log10 (P-value) |
|---|---|---|---|---|---|---|
| 12 | 9 | rs4883201 | TOTAL_CHOLESTROL | A2ML1 KLRG1 M6PR PHC1 | 1.742e-09 | 8.76 |
| 12 | 20 | rs11045163 | HDL | | 3.196e-09 | 8.5 |
| 12 | 21 | rs4149056 | TRIGS | RP11-125O5.2 SLCO1B1 | 1.782e-07 | 6.75 |
| 12 | 50 | rs10876041 | TOTAL_CHOLESTROL | DIP2B LARP4 | 1.028e-07 | 6.99 |
| 12 | 57 | rs3741414 rs11613352 | HDL TRIGS | AC126614.1 ARHGAP9 GLI1 INHBC INHBE MARS R3HDM2 RNU6-879P RP11-756H6.1 | 6.095e-14 9.397e-14 | 13.22 13.03 |
| 12 | 107 | rs10861661 | TRIGS | RFX4 RIC8B RP11-144F15.1 RP11-482D24.3 | 2.598e-07 | 6.59 |
| 12 | 109 | rs2241210 rs12321904 | HDL TOTAL_CHOLESTROL | KCTD10 MMAB UBE3B MVK RNU4-32P | 2.489e-20 2.077e-08 | 19.6 7.68 |
| 12 | 110 | rs10850380 rs10850380 | HDL TOTAL_CHOLESTROL | MMAB MVK RNU4-32P UBE3B | 4.058e-19 2.561e-08 | 18.39 7.59 |
| 12 | 111 | rs3184504 rs3184504 rs3184504 | HDL LDL TOTAL_CHOLESTROL | AC002395.1 ATXN2 RP3-473L9.4 SH2B3 | 4.097e-12 4.203e-12 1.623e-17 | 11.39 11.38 16.79 |
| 12 | 112 | rs653178 rs11065987 rs653178 | HDL LDL TOTAL_CHOLESTROL | ATXN2 RP11-686G8.1 RP11-686G8.2 U7 BRAP PCNPP1 | 1.064e-12 1.204e-11 1.038e-16 | 11.97 10.92 15.98 |
| 12 | 113 | rs12580178 | LDL | RPH3A | 2.307e-08 | 7.64 |
| 12 | 121 | rs1169288 rs2244608 | LDL TOTAL_CHOLESTROL | AC079602.1 C12orf43 HNF1A HNF1A-AS1 OASL RP11-216P16.2 | 6.447e-21 9.618e-18 | 20.19 17.02 |
| 12 | 122 | rs11057405 rs895953 | HDL TRIGS | CLIP1 RP11-512M8.3 RP11-512M8.5 VPS33A AC084018.1 HPD RHOF RP11-347I19.8 RP11-7M8.2 SETD1B TMEM120B | 1.231e-09 8.578e-08 | 8.91 7.07 |
| 12 | 123 | rs2454722 rs10773003 | HDL TOTAL_CHOLESTROL | HCAR1 HCAR2 HCAR3 RP11-324E6.6 RP11-324E6.9 C12orf65 CDK2AP1 MPHOSPH9 RNA5SP375 RP11-282O18.3 RP11-282O18.6 RP11-282O18.7 SBNO1 | 3.308e-14 4.083e-09 | 13.48 8.39 |
| 12 | 124 | rs7973683 rs11057408 | HDL TRIGS | CCDC92 DNAH10 DNAH10OS FAM101A RP11-214K3.18 RP11-214K3.19 RP11-214K3.20 RP11-214K3.21 RP11-214K3.22 RP11-214K3.23 RP11-214K3.5 RP11-380L11.3 RP11-380L11.4 ZNF664 | 5.259e-14 2.049e-12 | 13.28 11.69 |
| 12 | 125 | rs838876 | HDL | SCARB1 | 7.325e-33 | 32.14 |
| 12 | 133 | rs12423664 | TRIGS | FBRSL1 MUC8 RP11-503G7.1 RP11-503G7.2 | 8.189e-08 | 7.09 |

**Supplementary Table 15:** PheGWAS findings showcasing Chromosome Number, Marker Name, Associated Traits, Genes and the respective p-values for Chromosome 13

| Chr | Position group (Mb) | Marker Name | Associated Traits | Genes | P-Values | -log10 (P-value) |
|---|---|---|---|---|---|---|
| 13 | 32 | rs4942486 rs4942486 | LDL TOTAL_CHOLESTROL | BRCA2 IFIT1P1 N4BP2L1 RP11-298P3.4 SNORA16 | 2.261e-11 7.084e-08 | 10.65 7.15 |

**Supplementary Table 16:** PheGWAS findings showcasing Chromosome Number, Marker Name, Associated Traits, Genes and the respective p-values for Chromosome 14

| Chr | Position group (Mb) | Marker Name | Associated Traits | Genes | P-Values | -log10 (P-value) |
|---|---|---|---|---|---|---|
| 14 | 24 | rs8017377 rs6573778 | LDL TOTAL_CHOLESTROL | CBLN3 KHNYN NFATC4 NYNRIN SDR39U1 | 2.516e-15 2.958e-11 | 14.6 10.53 |
| 14 | 105 | rs4983559 | HDL | AKT1 CTD-3051D23.1 CTD-3051D23.4 LINC00638 RP11-982M15.2 RPS26P49 RPS2P4 SIVA1 ZBTB42 | 9.565e-09 | 8.02 |

**Supplementary Table 17:** PheGWAS findings showcasing Chromosome Number, Marker Name, Associated Traits, Genes and the respective p-values for Chromosome 15

| Chr | Position group (Mb) | Marker Name | Associated Traits | Genes | P-Values | -log10 (P-value) |
|---|---|---|---|---|---|---|
| 15 | 41 | rs721772 | HDL | ITPKA LTK RPAP1 TCEB1P2 TYRO3 | 2.358e-07 | 6.63 |
| 15 | 42 | rs2412710 rs2412710 | HDL TRIGS | CAPN3 GANC RNU6-188P RP11-164J13.1 ZNF106 | 1.36e-09 1.658e-11 | 8.87 10.78 |
| 15 | 43 | rs486301 | TRIGS | CKMT1B MAP1A PPIP5K1 TP53BP1 | 5.816e-08 | 7.24 |
| 15 | 44 | rs492571 rs16948098 | HDL TRIGS | FRMD5 PIN4P1 | 1.265e-12 4.838e-17 | 11.9 16.32 |
| 15 | 58 | rs10468017 rs588136 rs10468017 | HDL TRIGS TOTAL_CHOLESTROL | ALDH1A2 LIPC RP11-355N15.1 | 1.21e-188 3.365e-30 7.232e-48 | 187.92 29.47 47.14 |
| 15 | 59 | rs424346 | HDL | ADAM10 RN7SKP95 RP11-123C21.1 RP11-123C21.2 RP11-30K9.6 RP11-30K9.7 U3 | 4.84e-08 | 7.32 |
| 15 | 63 | rs2652834 rs2652834 | HDL TRIGS | LACTB RP11-244F12.2 RP11-69G7.1 RPS27L TPM1 | 3.592e-11 1.918e-08 | 10.44 7.72 |
| 15 | 72 | rs1035744 | TRIGS | CELF6 PARP6 PKM RP11-106M3.2 RP11-106M3.3 | 1.448e-07 | 6.84 |
| 15 | 73 | rs2415168 | TRIGS | ADPGK ADPGK-AS1 | 2.775e-07 | 6.56 |

**Supplementary Table 18:** PheGWAS findings showcasing Chromosome Number, Marker Name, Associated Traits, Genes and the respective p-values for Chromosome 16

| Chr | Position group (Mb) | Marker Name | Associated Traits | Genes | P-Values | -log10 (P-value) |
|---|---|---|---|---|---|---|
| 16 | 15 | rs3198697 | TRIGS | MIR1972-1 NTAN1 PDXDC1 RP11-680G24.4 RP11-680G24.5 RRN3 | 2.207e-08 | 7.66 |
| 16 | 30 | rs11649653 | TRIGS | AC106782.20 AC135048.13 BCL7C CTF1 CTF2P FBXL19 FBXL19-AS1 MIR4519 MIR762 ORAI3 | 1.56e-07 | 6.81 |
| 16 | 31 | rs749671 | TRIGS | AC135050.1 AC135050.5 BCKDK KAT8 PRSS53 RP11-196G11.1 RP11-196G11.2 RP11-196G11.4 STX4 VKORC1 ZNF646 ZNF668 | 6.106e-10 | 9.21 |
| 16 | 53 | rs1121980 rs9930333 | HDL TRIGS | FTO | 6.791e-09 3.251e-08 | 8.17 7.49 |
| 16 | 56 | rs9989419 rs247616 rs1800775 rs247616 | HDL LDL TRIGS TOTALCHOLESTROL | AC012181.1 CETP HERPUD1 NLRC5 RP11-325K4.2 RP11-325K4.3 RPS24P17 SLC12A3 | 0 2.566e-37 1.332e-26 4.47e-32 | 329.32 36.59 25.88 31.35 |
| 16 | 57 | rs59207181 rs1532624 rs7205804 rs1532624 | HDL LDL TRIGS TOTALCHOLESTROL | AC012181.1 CETP HERPUD1 NLRC5 RP11-322D14.1 RP11-325K4.2 RP11-325K4.3 | 0 1.844e-26 3.431e-25 1.39e-29 | 329.32 25.73 24.46 28.86 |
| 16 | 67 | rs16942887 rs3809630 | HDL TOTALCHOLESTROL | AC040162.1 CENPT CTC-479C5.10 CTC-479C5.11 CTC-479C5.12 CTC-479C5.17 CTRL EDC4 LCAT NRN1L NUTF2 PSKH1 PSMB10 SLC12A4 THAP11 RANBP10 TSNAXIP1 | 8.281e-54 1.544e-08 | 53.08 7.81 |
| 16 | 68 | rs16957696 rs13816 | HDL TOTALCHOLESTROL | AC130462.1 CTC-479C5.6 DDX28 DPEP2 DPEP3 DUS2 KARSP3 RNU6-359P SLC12A4 | 1.31e-46 8.945e-08 | 45.88 7.05 |
| 16 | 71 | rs8053891 rs8053891 | LDL TOTALCHOLESTROL | ATP5A1P3 DHODH IST1 PKD1L3 RP11-498D10.5 RP11-498D10.6 RP11-498D10.8 RPL39P31 | 3.76e-18 2.628e-18 | 17.43 17.58 |
| 16 | 72 | rs2000999 rs2000999 | LDL TOTALCHOLESTROL | DHODH DHX38 HP HPR PMFBP1 RP11-384M15.3 TXNL4B | 4.219e-41 6.804e-41 | 40.38 40.17 |
| 16 | 81 | rs2925979 rs2925979 | HDL TRIGS | CMIP RP11-391L3.5 | 1.321e-19 2.136e-07 | 18.88 6.67 |

**Supplementary Table 19:** PheGWAS findings showcasing Chromosome Number, Marker Name, Associated Traits, Genes and the respective p-values for Chromosome 17

| Chr | Position group (Mb) | Marker Name | Associated Traits | Genes | P-Values | -log10 (P-value) |
|---|---|---|---|---|---|---|
| 17 | 4 | rs8069974 | TOTAL_CHOLESTROL | ARRB2 CXCL16 GLTPD2 MED11 PLD2 PSMB6 RP11-314A20.5 RP11-81A22.5 TM4SF5 VMO1 ZMYND15 | 7.795e-08 | 7.11 |
| 17 | 7 | rs314253 rs314253 | LDL TOTAL_CHOLESTROL | ACADVL ASGR1 DLG4 DVL2 MIR324 PHF23 RPL7AP64 | 3.436e-10 2.808e-10 | 9.46 9.55 |
| 17 | 8 | rs4791641 | LDL | AURKB CTC1 LINC00324 PFAS RANGRF RP11-849F2.10 RP11-849F2.8 RP11-849F2.9 SLC25A35 | 1.314e-07 | 6.88 |
| 17 | 29 | rs2854322 | TOTAL_CHOLESTROL | AK4P1 NF1 RAB11FIP4 | 2.898e-07 | 6.54 |
| 17 | 37 | rs113612868 rs9972882 | HDL TOTAL_CHOLESTROL | AC087491.2 ERBB2 PGAP3 PNMT PPP1R1B STARD3 TCAP NEUROD2 | 5.447e-20 4.158e-08 | 19.26 7.38 |
| 17 | 38 | rs8069176 rs12309 | HDL TOTAL_CHOLESTROL | GSDMB IKZF3 LRRC3C ORMDL3 RP11-387H17.4 ZPBP2 CSF3 GSDMA MED24 PSMD3 RP11-387H17.6 SNORD124 THRA | 5.93e-13 1.953e-07 | 12.23 6.71 |
| 17 | 41 | rs231492 rs8077889 | HDL TRIGS | CD300LG FAM215A MPP2 PPY RP11-527L4.2 RP11-527L4.5 RP11-527L4.6 C17orf105 DUSP3 MPP3 RP5-905N1.2 SOST | 2.002e-07 9.879e-09 | 6.7 8.01 |
| 17 | 45 | rs6504872 rs6504872 | LDL TOTAL_CHOLESTROL | CTD-2026D20.2 EFCAB13 ITGB3 RP11-290H9.4 | 3.479e-13 6.987e-12 | 12.46 11.16 |
| 17 | 64 | rs1801689 | LDL | APOH CEP112 PSMD7P1 | 9.809e-12 | 11.01 |
| 17 | 65 | rs12602912 | TRIGS | AC006534.2 BPTF | 1.291e-07 | 6.89 |
| 17 | 66 | rs4148005 | HDL | ABCA8 | 5.743e-14 | 13.24 |
| 17 | 67 | rs2886232 rs2886232 | LDL TOTAL_CHOLESTROL | ABCA10 ABCA6 | 3.876e-11 3.874e-08 | 10.41 7.41 |
| 17 | 76 | rs4969178 | HDL | AC061992.1 AC061992.2 DNAH17 PGS1 RN7SL236P RP11-806H10.4 SNORA30 SOCS3 | 1.532e-12 | 11.81 |

**Supplementary Table 20:** PheGWAS findings showcasing Chromosome Number, Marker Name, Associated Traits, Genes and the respective p-values for Chromosome 18

| Chr | Position group (Mb) | Marker Name | Associated Traits | Genes | P-Values | -log10 (P-value) |
|---|---|---|---|---|---|---|
| 18 | 46 | rs2156498 | HDL | C18orf32 DYM MIR1539 RP11-110H1.1 RP11-110H1.4 RP11-110H1.9 RPL17 RPL17-C18orf32 SNORD58A SNORD58B SNORD58C SRP72P1 | 2.175e-07 | 6.66 |
| 18 | 47 | rs4939883 rs2156552 | HDL TOTAL_CHOLESTROL | LIPG RP11-813F20.1 | 1.796e-66 1.247e-31 | 65.75 30.9 |
| 18 | 57 | rs6567160 | HDL | RNU4-17P RP11-795H16.2 RP11-795H16.3 RPS3AP49 | 2.918e-09 | 8.53 |

**Supplementary Table 21:** PheGWAS findings showcasing Chromosome Number, Marker Name, Associated Traits, Genes and the respective p-values for Chromosome 19

| Chr | Position group (Mb) | Marker Name | Associated Traits | Genes | P-Values | -log10 (P-value) |
|---|---|---|---|---|---|---|
| 19 | 7 | rs4804833 rs7248104 | HDL TRIGS | AC010336.1 CTD-3193O13.1 CTD-3193O13.10 CTD-3193O13.11 CTD-3193O13.13 CTD-3193O13.8 CTD-3193O13.9 CTXN1 EVI5L LRRC8E MAP2K7 RN7SL115P RNA5SP463 SNAPC2 TGFBR3L TIMM44 INSR | 9.891e-08 5.045e-10 | 7 9.3 |
| 19 | 8 | rs2278236 rs4804311 | HDL TRIGS | AC010323.1 ANGPTL4 CTD-2550O8.7 KANK3 MARCH2 MIR4999 NDUFA7 RAB11B RAB11B-AS1 RPS28 AC092316.1 AC092316.2 AC130469.2 ADAMTS10 MYO1F PRAM1 ZNF414 | 3.185e-18 1.485e-09 | 17.5 8.83 |
| 19 | 10 | rs11881156 rs11881156 | LDL TOTALCHOLESTROL | AC007229.3 C19orf38 CARM1 DNM2 MIR199A1 TMED1 | 1.701e-55 8.708e-44 | 54.77 43.06 |
| 19 | 11 | rs737337 rs6511720 rs6511720 | HDL LDL TOTALCHOLESTROL | C19orf80 CTC-510F12.2 DOCK6 KANK2 RN7SL298P LDLR SMARCA4 SPC24 | 4.564e-17 3.85e-262 5.43e-202 | 16.34 261.42 201.26 |
| 19 | 18 | rs4808802 | TOTALCHOLESTROL | AC010335.1 CTD-3137H5.1 ELL ISYNA1 SSBP4 | 3.266e-08 | 7.49 |
| 19 | 19 | rs10401969 rs10401969 rs10401969 | LDL TRIGS TOTALCHOLESTROL | AC138430.4 HAPLN4 MAU2 NCAN SUGP1 TM6SF2 | 2.654e-54 9.702e-70 4.126e-77 | 53.58 69.01 76.38 |
| 19 | 33 | rs731839 rs731839 | HDL TRIGS | AKR1B1P7 CEBPG PEPD | 3.441e-09 2.651e-09 | 8.46 8.58 |
| 19 | 35 | rs1688030 | TRIGS | AC020907.1 CTD-2527I21.14 CTD-2527I21.9 FXYD3 GRAMD1A HPN HPN-AS1 SCN1B | 1.99e-07 | 6.7 |
| 19 | 44 | rs926054 rs926054 | LDL TOTALCHOLESTROL | AC069278.4 CTC-512J12.4 CTC-512J12.6 ZNF180 ZNF229 ZNF285 ZNF285B | 3.358e-15 2.407e-08 | 14.47 7.62 |
| 19 | 45 | rs2075650 rs7254892 rs439401 rs7412 | HDL LDL TRIGS TOTALCHOLESTROL | APOC1 APOC1P1 APOC4 APOC4-APOC2 APOE CTB-129P6.4 PVRL2 TOMM40 APOC2 CLPTM1 CTB-129P6.11 | 9.716e-26 0 1.423e-66 1.56e-283 | 25.01 329.32 65.85 282.81 |
| 19 | 46 | rs7255743 rs7255743 | LDL TOTALCHOLESTROL | ERCC1 FOSB OPA3 PPM1N RTN2 VASP | 1.408e-25 9.61e-13 | 24.85 12.02 |
| 19 | 49 | rs492602 rs516246 | LDL TOTALCHOLESTROL | FUT1 FUT2 IZUMO1 MAMSTR NTN5 RASIP1 RN7SL345P SEC1P | 9.422e-14 9.133e-17 | 13.03 16.04 |
| 19 | 50 | rs1132990 | TRIGS | CTD-3148I10.13 CTD-3148I10.15 CTD-3148I10.9 FCGRT FLT3LG MIR150 NOSIP RCN3 RPL13A RPS11 SNORD32A SNORD33 SNORD34 SNORD35A SNORD35B Y_RNA hsa-mir-150 | 6.125e-08 | 7.21 |
| 19 | 52 | rs17695224 | HDL | AC006272.1 AC006272.2 FPR1 FPR3 ZNF577 | 2.417e-13 | 12.62 |
| 19 | 54 | rs103294 rs103294 | HDL TOTALCHOLESTROL | AC008984.2 AC008984.5 AC008984.6 AC008984.7 AC010492.4 AC010492.5 AC010518.3 AC098789.1 CTD-2337J16.1 LILRA3 LILRA4 LILRA5 LILRB2 LILRB5 MIR4752 RNU6-1307P RPS9 VN1R104P | 3.995e-30 2.27e-11 | 29.4 10.64 |

**Supplementary Table 22:** PheGWAS findings showcasing Chromosome Number, Marker Name, Associated Traits, Genes and the respective p-values for Chromosome 20

| Chr | Position group (Mb) | Marker Name | Associated Traits | Genes | P-Values | -log10 (P-value) |
|---|---|---|---|---|---|---|
| 20 | 12 | rs364585 | LDL | RP11-157E14.1 RP5-1069C8.2 RP5-1069C8.3 SPTLC3 | 4.278e-10 | 9.37 |
| 20 | 17 | rs2328223 | LDL | AL035045.1 RNU6-192P RP5-905G11.3 | 5.632e-09 | 8.25 |
| 20 | 33 | rs6058202 | HDL | AL121753.1 EDEM2 MMP24 MMP24-AS1 MT1P3 PROCR RNA5SP483 RP4-614O4.12 | 1.275e-07 | 6.89 |
| 20 | 34 | rs7264396 rs7264396 | LDL TOTAL_CHOLESTROL | C20orf173 ERGIC3 FER1L4 RP3-477O4.16 RP3-477O4.5 RPL36P4 RPL37P1 SPAG4 | 4.412e-08 6.585e-13 | 7.36 12.18 |
| 20 | 39 | rs6065311 rs6029143 rs2235367 | LDL TRIGS TOTAL_CHOLESTROL | PLCG1 RP1-1J6.2 TOP1 RP3-511B24.5 RP3-511B24.6 RPL23AP81 ZHX3 | 1.656e-30 4.933e-08 7.219e-25 | 29.78 7.31 24.14 |
| 20 | 40 | rs4142393 rs4142393 | LDL TOTAL_CHOLESTROL | CHD6 RP4-620E11.4 | 1.347e-09 2.743e-07 | 8.87 6.56 |
| 20 | 43 | rs1800961 rs1800961 rs1800961 | HDL LDL TOTAL_CHOLESTROL | AL132772.1 C20orf62 HNF4A MIR3646 RP5-1013A22.2 RP5-1013A22.5 RP5-881L22.4 RP5-881L22.6 | 1.639e-34 6.034e-10 1.342e-24 | 33.79 9.22 23.87 |
| 20 | 44 | rs4465830 rs4810479 | HDL TRIGS | FTLP1 PCIF1 PLTP ZNF335 CTSA NEURL2 RP3-337O18.9 SPATA25 ZSWIM1 ZSWIM3 | 5.175e-40 2.067e-34 | 39.29 33.68 |
| 20 | 45 | rs6066141 | TRIGS | EYA2 GAPDHP54 | 2.339e-08 | 7.63 |

**Supplementary Table 23:** PheGWAS findings showcasing Chromosome Number, Marker Name, Associated Traits, Genes and the respective p-values for Chromosome 22

| Chr | Position group (Mb) | Marker Name | Associated Traits | Genes | P-Values | -log10 (P-value) |
|---|---|---|---|---|---|---|
| 22 | 21 | rs113359481 rs111562164 | HDL TOTAL_CHOLESTROL | RIMBP3C RN7SKP221 SCARNA17 SCARNA18 UBE2L3 CCDC116 YDJC | 4.303e-18 4.363e-10 | 17.37 9.36 |
| 22 | 30 | rs5763662 rs5763662 | LDL TOTAL_CHOLESTROL | AC003681.1 CTA-85E5.10 MTMR3 | 1.191e-08 7.719e-08 | 7.92 7.11 |
| 22 | 35 | rs138777 | TOTAL_CHOLESTROL | HMGXB4 MIR3909 RP3-510H16.3 TOM1 | 4.74e-08 | 7.32 |
| 22 | 38 | rs3761445 | TRIGS | AL021977.1 MAFF PLA2G6 RN7SL704P RP1-5O6.5 TMEM184B | 8.062e-12 | 11.09 |
| 22 | 46 | rs4253776 rs4253772 | LDL TOTAL_CHOLESTROL | CDPF1 PKDREJ PPARA TTC38 | 3.353e-08 9.852e-09 | 7.47 8.01 |

#### FIGURES:

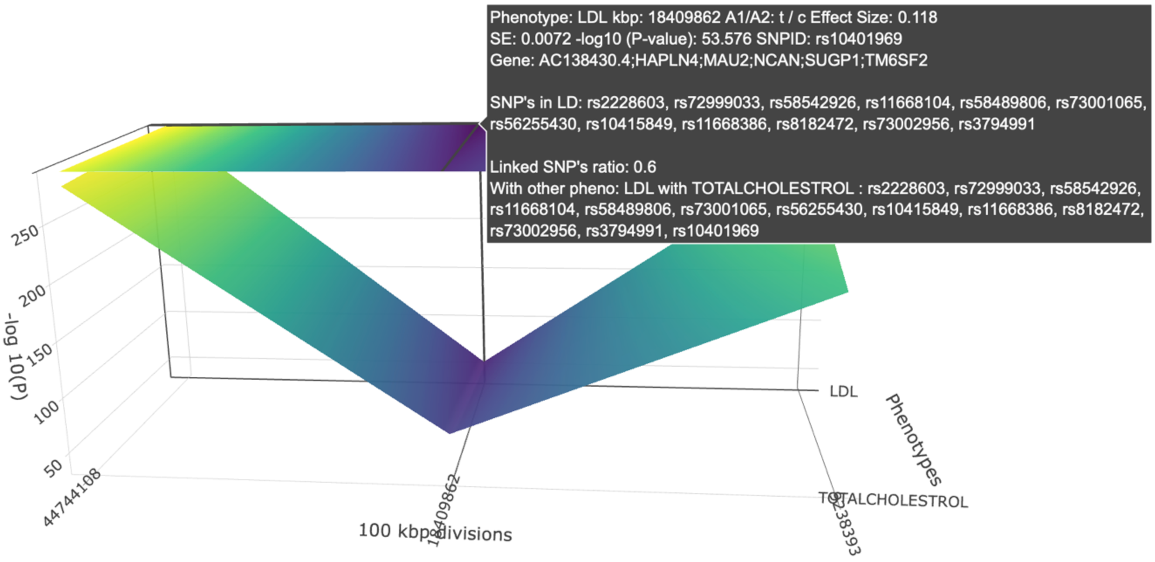


*Fig 1: PheGWAS diagram for chromosome 19 on Total Cholesterol and LDL.*

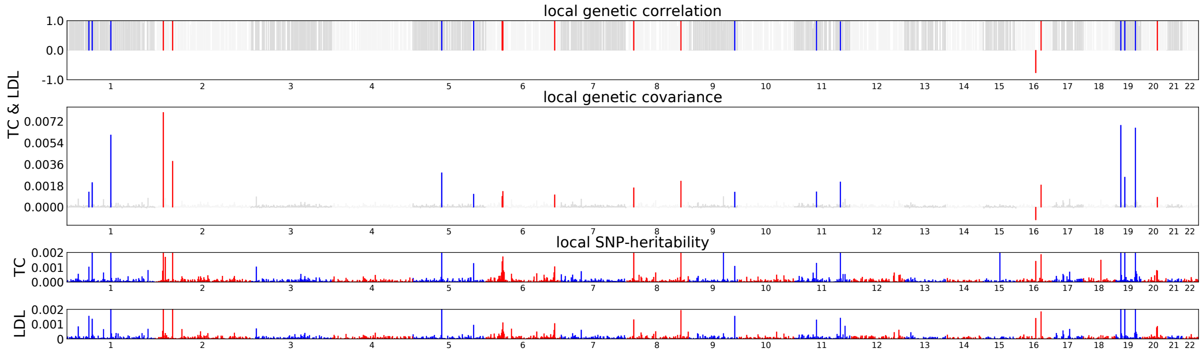


*Fig 2: Estimates of Local Genetic Covariance for the Pairs of Traits LDL & TC using ρ-HESS. In 19th chromosome it shows 3 peaks for local genetic correlation.*

**Section 1: Colocalization analysis performed for chromosome 19 for BP region from 44744108 to 46102697.**

**Section 1a:** Using HyprColoc

For each iteration of the algorithm, HyPrColoc returns:

i. a cluster of putatively colocalized traits
ii. the posterior probability that these traits are colocalized
iii. the 'regional association' probability* (which is always > the posterior probability)
iv. a candidate causal variant explaining the shared association
v. the proportion of the posterior probability explained by this variant (which represents the HyPrColoc multi-trait fine-mapping probability).

Following is the output:

```
Call:
hyprcoloc

Results:
  iteration                         traits posterior_prob regional_prob candidate_snp posterior_explained_by_snp dropped_trait
1         1 LDL_beta, TOTALCHOLESTROL_beta              1             1        rs7412                          1          <NA>
2         2                           None             NA             1          <NA>                         NA      HDL_beta
```

**Section 1b:** Using MOLOC

The output is a list with three elements:

1. First element is a data frame containing the priors, likelihoods and Posteriors for each locus and each combination. We usually care about the last columns, the posterior probability of a shared variant for each model;
2. Second element is the number of SNPs analyzed, in common across all datasets;
3. Third element of the output is a data frame with the posterior probability of the most likely SNP.

Following is the output.

```
> print(moloc[[1]])
          prior sumbf logBF_locus     PPA
a         1e-04    47          43 0.0e+00
a,b       1e-08  1746        1737 0.0e+00
a,c       1e-08   799         789 0.0e+00
a,d       1e-08   136         126 0.0e+00
a,bc      1e-09  2499        2489 3.3e-38
a,bd      1e-09  1815        1805 0.0e+00
a,cd      1e-09   867         857 0.0e+00
a,bcd     1e-10  2568        2558 1.5e-09
a,b,c     1e-12  2498        2488 8.4e-42
a,b,d     1e-12  1835        1826 0.0e+00
a,b,cd    1e-13  2566        2556 3.9e-13
a,c,d     1e-12   888         878 0.0e+00
a,bd,c    1e-13  2565        2555 7.8e-14
a,bc,d    1e-13  2588        2579 1.7e-03
a,b,c,d   1e-16  2588        2578 1.2e-06
b         1e-04  1700        1695 0.0e+00
b,c       1e-08  2452        2442 6.2e-58
b,d       1e-08  1788        1778 0.0e+00
ac,b      1e-09  2498        2488 1.1e-38
ad,b      1e-09  1816        1806 0.0e+00
b,cd      1e-09  2520        2510 2.7e-29
acd,b     1e-10  2566        2557 5.1e-10
b,c,d     1e-12  2541        2531 2.9e-23
ad,b,c    1e-13  2568        2558 3.0e-12
ac,b,d    1e-13  2588        2578 7.6e-04
c         1e-04   753         748 0.0e+00
c,d       1e-08   841         831 0.0e+00
ab,c      1e-09  2499        2489 1.7e-38
ad,c      1e-09   869         859 0.0e+00
bd,c      1e-09  2520        2510 3.8e-29
abd,c     1e-10  2567        2557 1.1e-09
ab,c,d    1e-13  2587        2578 6.6e-04
d         1e-04    88          83 0.0e+00
ab,d      1e-09  1835        1825 0.0e+00
ac,d      1e-09   888         878 0.0e+00
bc,d      1e-09  2541        2531 4.2e-20
abc,d     1e-10  2588        2578 1.0e+00
ab        1e-05  1747        1742 0.0e+00
ab,cd     1e-10  2567        2557 1.1e-09
ac        1e-05   800         795 0.0e+00
ac,bd     1e-10  2567        2557 1.1e-09
ad        1e-05   117         112 0.0e+00
ad,bc     1e-10  2569        2559 5.6e-09
bc        1e-05  2453        2448 2.0e-54
bd        1e-05  1768        1763 0.0e+00
cd        1e-05   821         816 0.0e+00
abc       1e-06  2500        2495 4.8e-35
abd       1e-06  1815        1810 0.0e+00
acd       1e-06   868         863 0.0e+00
bcd       1e-06  2521        2516 1.1e-25
abcd      1e-07  2568        2563 3.0e-06
zero      1e+00     0           0 0.0e+00
```

```
> print(moloc[[2]])
[1] 137
> print(moloc[[3]])
     coloc_ppas best.snp.coloc
a       1.0e+00         rs7412
b       1.0e+00         rs7412
ab      1.0e+00         rs7412
c       1.0e+00         rs7412
ac      7.7e-04         rs7412
bc      1.0e+00         rs7412
abc     1.0e+00         rs7412
d       3.1e-06         rs7412
ad      3.0e-06         rs7412
bd      3.0e-06         rs7412
abd     3.0e-06         rs7412
cd      3.0e-06         rs7412
acd     3.0e-06         rs7412
bcd     3.0e-06         rs7412
abcd    3.0e-06         rs7412
```
